# *Phlebotomus papatasi* sand fly salivary protein diversity and immune response potential in Egypt and Jordan populations

**DOI:** 10.1101/649517

**Authors:** Catherine M. Flanley, Marcelo Ramalho-Ortigao, Iliano V. Coutinho-Abreu, Rami Mukbel, Hanafi A. Hanafi, Shabaan S. El-Hossary, Emadeldin Y. Fawaz, David F. Hoel, Alexander W. Bray, Gwen Stayback, Douglas A. Shoue, Shaden Kamhawi, Scott Emrich, Mary Ann McDowell

## Abstract

*Phlebotomus papatasi* sand flies inject their hosts with a myriad of pharmacologically active salivary proteins to assist with blood feeding and to modulate host defenses. These salivary proteins have been studied for their role in cutaneous leishmaniasis disease outcome with different salivary proteins attenuating or exacerbating lesion size. Studies have shown that while co-administered sand fly saliva exacerbates *Leishmania major* infections in naïve mice, animals pre-exposed to saliva are protected, with the infection attenuated via a delayed-type hypersensitivity immune reaction. These studies highlight the potential of the salivary components to be used as a vaccine. One protein in particular, *P. papatasi* salivary protein 15 (PpSP15) has been intensively studied because of its ability to protect mice against *Le. major* challenge. The number of antigenic molecules included in vaccines is restricted thus emphasizing the role of population genetics to identify molecules, like PpSP15, that are functionally significant, conserved across populations and do not experience selection. Three distinct ecotope study sites, one in Egypt (Aswan) and two in Jordan (Swaimeh and Malka), were chosen based on their elevation, rainfall, vegetation, differing reservoir species, and the presence or absence of *Le. major*. The objective of this work was to analyze the genetic variability of nine of the most abundantly expressed salivary proteins including PpSP12, PpSP14, PpSP28, PpSP29, PpSP30, PpSP32, PpSP36, PpSP42, and PpSP44 and to predict their ability to elicit an immune response. Two proteins, PpSP12 and PpSP14, demonstrated low genetic variability across the three sand fly populations represented in this study, with multiple predicted MHCII epitope binding sites, identified by alleles present in the human populations from the study sites. The other seven salivary proteins revealed greater allelic variation across the same sand fly populations indicating that their use as vaccine targets may prove to be challenging.

## Author Summary

*Phlebotomus papatasi* sand flies vector *Leishmania major* parasites, one of the causative agents of cutaneous leishmaniasis (CL). Approximately 0.7-1.2 million cases of CL occur each year. CL produces disfiguring skin lesions for which no vaccine currently exists. Hematophagous vector salivary proteins are pharmacologically active molecules that modulate inflammation, vasoconstriction, and blood clotting for females that require a sanguineous meal for oviposition. Salivary proteins from multiple phlebotomine sand fly species have been widely studied and scrutinized to characterize their function in blood feeding facilitation as well as their ability to exacerbate or attenuate *Leishmania* infections and their potential as vaccine candidates. A successful sand fly salivary protein-based vaccine to combat CL largely depends on the genetic variability, expression profiles, and human immune response to the salivary proteins selected from geographically distant sand fly populations. The purpose of this study was to analyze these parameters in nine abundantly expressed *P. papatasi* salivary proteins from three distinct ecotopes in Egypt and Jordan in order to assess their potential as vaccine targets

## Introduction

Leishmaniasis is a group of neglected diseases caused by *Leishmania* parasites, vectored by phlebotomine sand flies and endemic in 98 countries [1]. Different *Leishmania* species can be or are uniquely associated with distinct clinical outcomes, ranging from cutaneous lesions to fatal visceral disease. Sand flies on the other hand may be specific or not for the transmission of *Leishmania* spp. *Phlebotomus papatasi* sand flies are specific vectors of *Leishmania major*, one of the causative agents of cutaneous leishmaniasis (CL) [2]. Approximately 0.7-1.2 million cases of CL occur each year [1]. CL produces scarring skin lesions and current treatments can be toxic, expensive, require multiple administrations, and can be difficult to access [2]. Although significant effort has been expended, there currently is no efficacious vaccine for human populations.

Salivary proteins of hematophagous insects are pharmacologically active molecules that modulate inflammation, vasoconstriction, and blood clotting [3]. In *P. papatasi* infected with *Le. major*, parasites are regurgitated into the host’s skin during probing or feeding, along with a cocktail of salivary proteins, where an infection can be established. Salivary proteins from various phlebotomine sand fly species have been characterized with regards to their function in blood feeding and their effectiveness as markers of exposure [4–6]. Sand fly salivary proteins can exacerbate or attenuate *Leishmania* infections [7–9], and have been suggested as potential as vaccine candidates [10,11].

It has been previously determined that exposure to uninfected *P. papatasi* bites confers some level of protection against *Le. major* in murine models, presumably via stimulation of a delayed-type hypersensitivity immune response at the site of inoculation [12]. Over thirty different salivary proteins are inoculated into a host with each *P. papatasi* bite [13]. One particular salivary protein, PpSP15, induces a Th-1 mediated immune response with a hallmark increase in IFN-γ in mice [8], and vaccination of nonhuman primates with the *P. duboscqi* orthologous PdSP15 resulted in a significant decrease in parasite load and lesion size, though full protection was not established [10]. Conversely, in naïve hosts, sand fly saliva exacerbates disease progression by downregulating the host’s immune response while polarizing the immune response to favor Th2 cytokine production [8,14–16].

Several questions remain concerning the role of sand fly salivary proteins in the epidemiology of leishmaniasis, particularly those pertaining to cross species protection and genetic variability. Cross-protective effects of saliva from different phlebotomine species is another current area of vaccine development research. A cross-protective effect against *Le. major* was demonstrated in mice pre-exposed to uninfected *P. papatasi* sand fly bites and subsequently challenged with *Le. major* plus either *P. papatasi* or *P. duboscqi* salivary gland homogenate [17]. Both pre-exposed groups revealed smaller lesion sizes and a decreased parasitic load compared to the unexposed controls [17]. A similar study was conducted with New World sand fly species where hamsters inoculated with *Lu. longipalpis* salivary gland homogenate or a DNA plasmid coding for the highly expressed *Lu. longipalpis* salivary protein LJM19, were protected against challenge with either *Le. braziliensis* and *Lu. longipalpis* salivary gland homogenate, as well as *Le. braziliensis* and *Lu. intermedia* salivary gland homogenate [18].

Immunodominant sand fly salivary proteins have also been exploited in epidemiological studies as markers of exposure and risk for *Leishmania* transmission. Anti-saliva antibodies correlate to intensity of exposure with higher anti-saliva antibody titers indicating greater exposure to sand fly bites and a greater probability of transmission [16,19,20]. A disadvantage of measuring antibodies against whole sand fly saliva is the possibility of cross-reactivity between different sand fly species as they may share a significant number of salivary protein antigens. Ideally, a single species-specific salivary protein could be identified to determine exposure and transmission risk (reviewed in [21]). PpSP32, an immunodominant 32 kDa protein present in *P. papatasi* saliva, has been validated as an exposure screening tool in Tunisia and Saudi Arabia with no cross-reactivity detected when tested against *P. perniciosus*, a vector of *Le. infantum*, which is also commonly found in sympatry with *P. papatasi* [4,6,22].

Genetic variability among populations of sand flies will influence the success of any salivary protein-based vaccine. Specifically, highly polymorphic salivary proteins and those under positive selection should be cautiously considered for further vaccine development. PpSP15, sampled from sand fly populations in Egypt, Jordan, Saudi Arabia, Israel, and Sudan demonstrated minimal selection with a high degree of conservation at the amino acid level validating PpSP15 as a potential vaccine candidate [23,24]. Here we analyzed the genetic variability of nine highly expressed *P. papatasi* salivary proteins including PpSP12, PpSP14, PpSP28, PpSP29, PpSP30, PpSP32, PpSP36, PpSP42, and PpSP44 from three representative *P. papatasi* populations from Egypt and Jordan.

As a geographically widespread species, *P. papatasi* is prevalent in the Mediterranean Basin, especially the Middle East and North Africa, with an ability to adapt to a variety of habitats that exhibit different climates, elevations, vegetation, and host species. As a result of adaptation to ecological variation, it is expected that sand flies and their salivary proteins face selective pressures that could influence vector competency and disease outcomes [25]. *P. papatasi* population genetics studies have demonstrated that although pockets of genetic variability exist between populations, evidence suggests that the species as a whole remains relatively homogeneous [26–31].

Even so, it remains vitally important to continually monitor salivary protein genetic variability to ensure the most appropriate salivary proteins are chosen as vaccine targets. A successful sand fly salivary protein-based vaccine to combat CL also depends on expression profiles and human (host) immune response to these salivary proteins, ideally selected from geographically distant sand fly populations. Egypt and Jordan are classified as endemic areas for CL, with certain regions designated as hyperendemic in Jordan, with *Le. major* causing the majority of CL cases but *Le. tropica* incriminated in Jordan as well [32,33]. The purpose of this study was to analyze 9 abundantly expressed *P. papatasi* salivary proteins as potential vaccine targets that are conserved across populations from distinct ecotopes in Egypt and Jordan and demonstrate the potential to elicit an immune response, similar to PpSP15 [24]. We recommend that PpSP12 and PpSP14 also be considered for further vaccine development as we show that they are conserved across populations while also have potential to elicit an immune response. We caution of the use of a highly variable protein like PpSP28 for further development.

## Methods

### Sand flies

*P. papatasi* were collected from one field site in each of the following locations: Aswan, Egypt (GPS coordinates N 24°10’, E 32°52’), Malka, Jordan (GPS coordinates 31°48’, E 35°35’), and Swaimeh, Jordan (N 32°40’, E 35°45’), in 2006 and 2007 (Fig 1). Both CO_2_ baited (Aswan) and non-baited (Malka and Swaimeh) CDC-style light traps collected sand flies between the hours of 18:00 and 06:00. Three trappings were attempted each year in 2006 and 2007: early (June), middle (August), and late (September). One collection occurred in Malka in late (September) 2006 while three collections occurred in Swaimeh and Aswan in late (September) 2006, early (June), and middle (August) 2007. Sand flies remained alive until dissection and were euthanized in soapy water. Flies were individually identified by microscopic examination of female spermathecae according to Lane [34], and only non-parous females were used in the analysis presented. Parity was assessed according to Anez [35].

**Fig 1.**
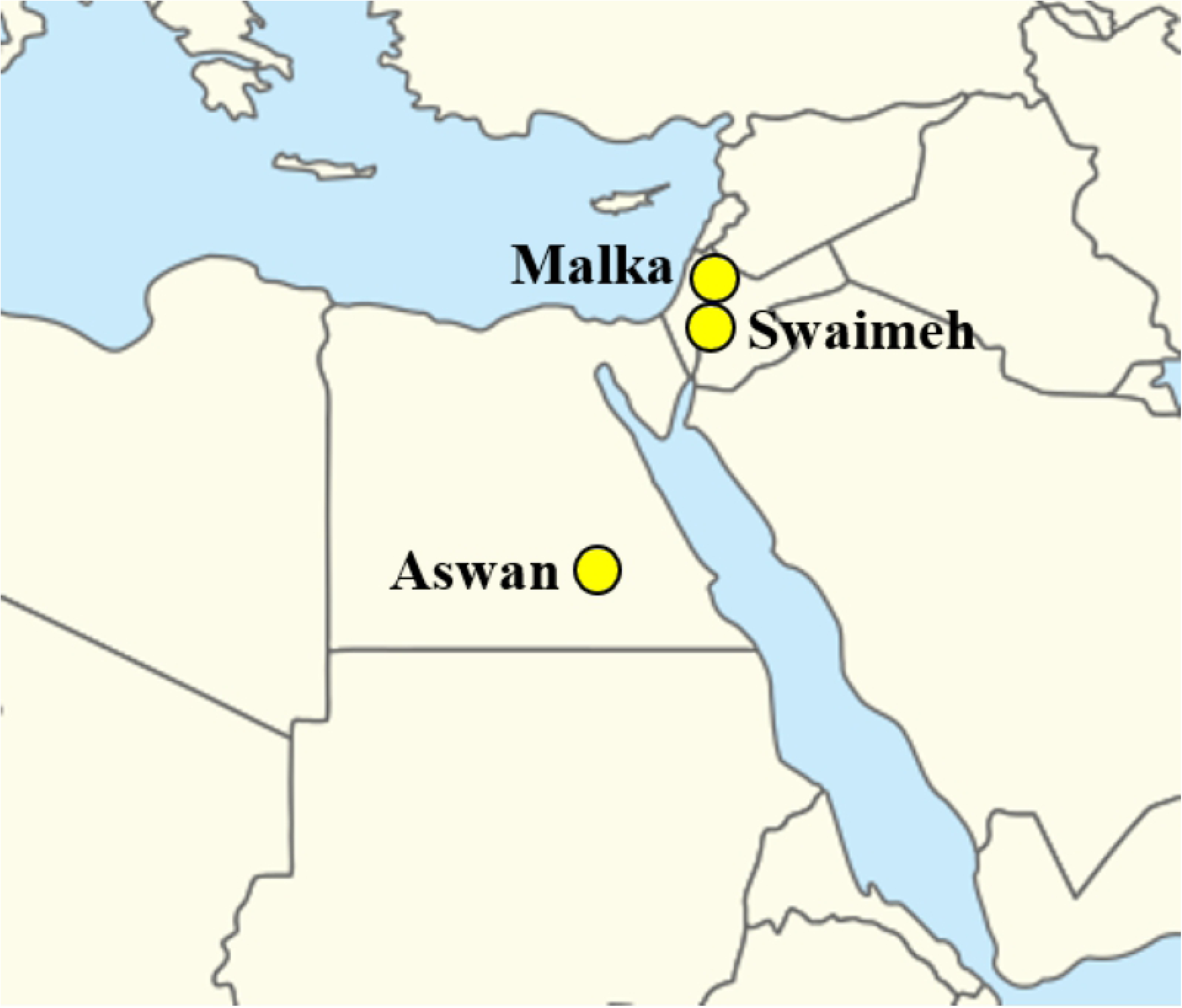
*Phlebotomus papatasi* study collection sites. Map adopted from Wikimedia Commons by Styx under public domain [36].

*P. papatasi* from Aswan (PPAW) were collected from a small village near the Nile River that permits artificial irrigation for the cultivation of crops like corn (*Zea mays*), wheat (*Triticum aestivum*), mangoes (*Mangifera indica*), and date palms (*Phoenix dactylifera*). Dogs, goats, and cattle are kept and raised in the village as well. The village sits at 117 m above sea level. Temperatures typically fall between 24° C and 45° C with minimal rainfall. Sand flies are abundant in this village though *Le. major* is absent [37]. *P. papatasi* collected from Swaimeh (PPJS) inhabit an area endemic for zoonotic *Le. major* due to the presence of *Psammomys obesus*, the reservoir host [38]. This low elevation area (∼350m below sea level) experiences a Saharan Mediterranean climate with rainfall less than 50mm that occurs from November to April. Temperatures maximally range from 35-40° C in summer months and minimally range from 8-12° C in the winter. The sandy, rocky, salty soil supports halophytic and tropical flora species such as chenopods [39]. *P. papatasi* collected from Malka (PPJM) inhabit a rocky landscape with a typical Mediterranean climate. Malka is located at an elevation of 670 m. During the collection time in 2006, only *Le. tropica* was present in the region and *Le. major* was absent hypothesized due to the absence of *Ps. obesus* [40].

### Sample preparation

Dissected, *P. papatasi* female heads with both salivary glands intact were placed in 1.5 ml centrifuge tubes with 50 μL RNA later (Ambion, Austin, TX, USA) and homogenized with an RNAse-free pestle and hand-held homogenizer. Samples were stored at 4° C for up to 48 hours, shipped on dry ice, and then stored at -80° C until analyzed. Table 1 outlines the number of individuals from each site for each salivary protein.

**Table 1.** ***Phlebotomus papatasi* salivary protein amplicon lengths and number of individual sand flies per collection site.**

| Salivary Protein | Amplicon length (bp) | All | PPAW | PPJM | PPJS |
| --- | --- | --- | --- | --- | --- |
| PpSP12 | 291 | 96 | 26 | 29 | 41 |
| PpSP14 | 246 | 119 | 29 | 44 | 46 |
| PpSP28 | 554 | 111 | 26 | 30 | 55 |
| PpSP29 | 651 | 126 | 46 | 38 | 42 |
| PpSP30 | 183 | 70 | 20 | 28 | 22 |
| PpSP32 | 568 | 130 | 42 | 45 | 43 |
| PpSP36 | 637 | 82 | 25 | 22 | 35 |
| PpSP42 | 614 | 109 | 27 | 45 | 37 |
| PpSP44 | 675 | 121 | 35 | 44 | 42 |

Amplicon lengths = base pairs. PPAW: Aswan, Egypt; PPJM: Malka, Jordan; PPJS: Swaimeh, Jordan.

### RNA extraction and cDNA synthesis

Total RNA was extracted from the heads and salivary glands of individual *P. papatasi* samples using the RNeasy Mini RNA isolation kit (Qiagen, Valencia, CA, USA). cDNAs were synthesized using Invitrogen reagents (Invitrogen, Carlsbad, CA, USA), per manufacturer’s specifications and briefly outlined in Coutinho-Abreu et al 2011 [41] and Ramalho-Ortigão et al 2015 [24].

### Sequence analyses

cDNAs produced from the total RNA of individual *P. papatasi* were amplified by PCR. Primers used to amplify each salivary protein can be found in S1 Table. PCR products were purified by twice washing in 150 μL DNAse/RNAse-free water (Invitrogen, Carlsbad, CA, USA) in Multiscreen PCR cleaning plates (Millipore, Burlington, Massachusetts, USA) with vacuum application (10 psi). Purified PCR products were resuspended in 50 μL sterile water. Leading and lagging strands were sequenced and poor-quality sequences were excluded from the analyses. Forward and reverse chromatograms were inspected and consensus sequences were aligned using MEGA [42] and manually corrected. The resulting sequences were deposited in GenBank (http://ncbi.nlm.nih.gov) and accession numbers can be found in S1 Table.

### Multi-copy assessment of salivary proteins

We looked for copy number variation relative to currently assembled *P. papatasi* loci by following the approach of Miles *et al*. [43]. In short, paired sequences from the two individual entries with the most reads (SRR1997534 and SRR199776) were downloaded from SRA. After initial overall quality checking using FastQC [44], these paired reads were then aligned to the Ppap reference assembly using BWA 0.5.9r16 [45]. Next, based on the resulting alignments (SAM output), reads were placed into non-overlapping 300 bp bins such that each bin contained all reads whose alignment started in its corresponding 300 bp interval. Unlike Miles *et al*. [43] there were no known “core genome” coordinates that excluded repeat regions for computing a less biased average count for normalization. Therefore, discrete counts in each bin were normalized based on the average count across all scaffolds, median count, and average count excluding terminal 2 kb of scaffolds (1 kb on 5’ and 3’ of scaffolds). Although normalization slightly differed, the results in terms of under (less than 0.5 average/median) and over (more than 2 average/median) remained the same within and between the two samples considered. Because of under assembly of heterologous regions, the final normalization value was computed based on the empirical distribution of read bin counts with the value with the greatest number of entries near the computed overall average. Significance was derived using a Poisson model parameterized with this estimate as lambda using the Lander-Waterman model of sequence sampling [46], and a Bonferroni correction was applied to correct for multiple comparisons.

### Population analyses

Both interpopulation and intrapopulation analyses were performed using DnaSP v.6 [47]. Interpopulation parameters assessed included: fixation indexes such as Fst [48,49] and Gst [50], as well as Hs and Ks indexes [51]. Other parameters assessed included: neutral evolution hypothesis [52] and neutrality tests Tajima’s D [53] and Fu and Li’s D and F [54]. The Ka/Ks ratio (ω) for the whole salivary protein as well as a sliding window analysis of 70 codons each was calculated.

Weblogos [55] pictorially depict the relative frequencies of polymorphic nucleotides and amino acids. The height of the bases indicate relative frequency and conservation is depicted by the overall weight of the stack. Network 5 [56] generated median joining networks exhibiting haplotype relationships.

### Secondary structure and T-cell epitope predictions

Secondary structure predictions for each salivary protein were generated using a secondary structure prediction tool (http://bioinf.cs.ucl.ac.uk/psipred) with default parameters based on the consensus sequence for all individual amino acid sequences from DnaSP. Two different predictions tools predicted the promiscuous HLA-class II binding sites and human T-cell epitopes: IEDB analysis resource T-cell epitope prediction tools (http://tools.immuneepitope.org/main/html/tcell_tools.html) [57,58] and ProPred MHC class II binding prediction server (http://www.imtech.res.in/raghava/propred/) [59]. For the 51 HLA alleles tested in ProPred, thresholds included a promiscuous search set to 3%. For the 27 HLA alleles tested in IEDB, only predicted peptides with a Consensus percentile rank of 0.10 or below are included as the top 10% of peptides with the strongest predicted binding affinity.

## Results

Using previously published PpSP15 [24] data as a guide to help highlight the proteins presented in this study, we would prioritize PpSP12 and PpSP14 for vaccine development according to our in-depth analyses presented below. The remaining seven salivary proteins may be valid vaccine components; however, the extent of the allelic variation present suggests that their development as vaccine components may prove challenging. Herein, we present data for PpSP28 as a representative salivary protein with PpSP29, PpSP30, PpSP32, PpSP36, PpSP42, and PpSP44 provided in the supplemental materials.

### PpSP12 in-depth analyses

#### Nucleotide and amino acid genetic diversity

The 291 bp *PpSP12* fragment produced 14 polymorphic sites (Fig 2A). Two specific nucleotide positions indicate limited heterogeneity between the Jordan populations compared to the Egypt population. Position one shows conservation of adenine in both Jordan populations with variation present in the Egypt population. Conversely, in position 13 the conserved frequency of adenine is greater in the Aswan population in comparison to both Malka and Swaimeh. All of the populations present similar heterogeneity at positions 6-9. Although heterogeneity exists in *PpSP12*, it is the lowest when comparing all 9 salivary proteins. The translated PpSP12 amino acid sequence has 6 variable positions out of 97 total amino acids (Fig 2B). At position 2, arginine and lysine are both found in all populations. Both arginine and lysine are positively-charged and belong to the basic group of amino acids. They are frequently substituted for each other in nature [60]. The frequency of alanine and proline are relatively equal for all populations in position 4. Both of these amino acids are small in size, nonpolar, and hydrophobic. At position 3 and 5, the relative frequencies of lysine and asparagine are the same except the Aswan population at position 5 has a much higher frequency of lysine. Lysine is a positively charged polar amino acid and asparagine is polar but neutrally charged. Both are frequently found in protein active, or binding, sites [60].

**Fig 2.**
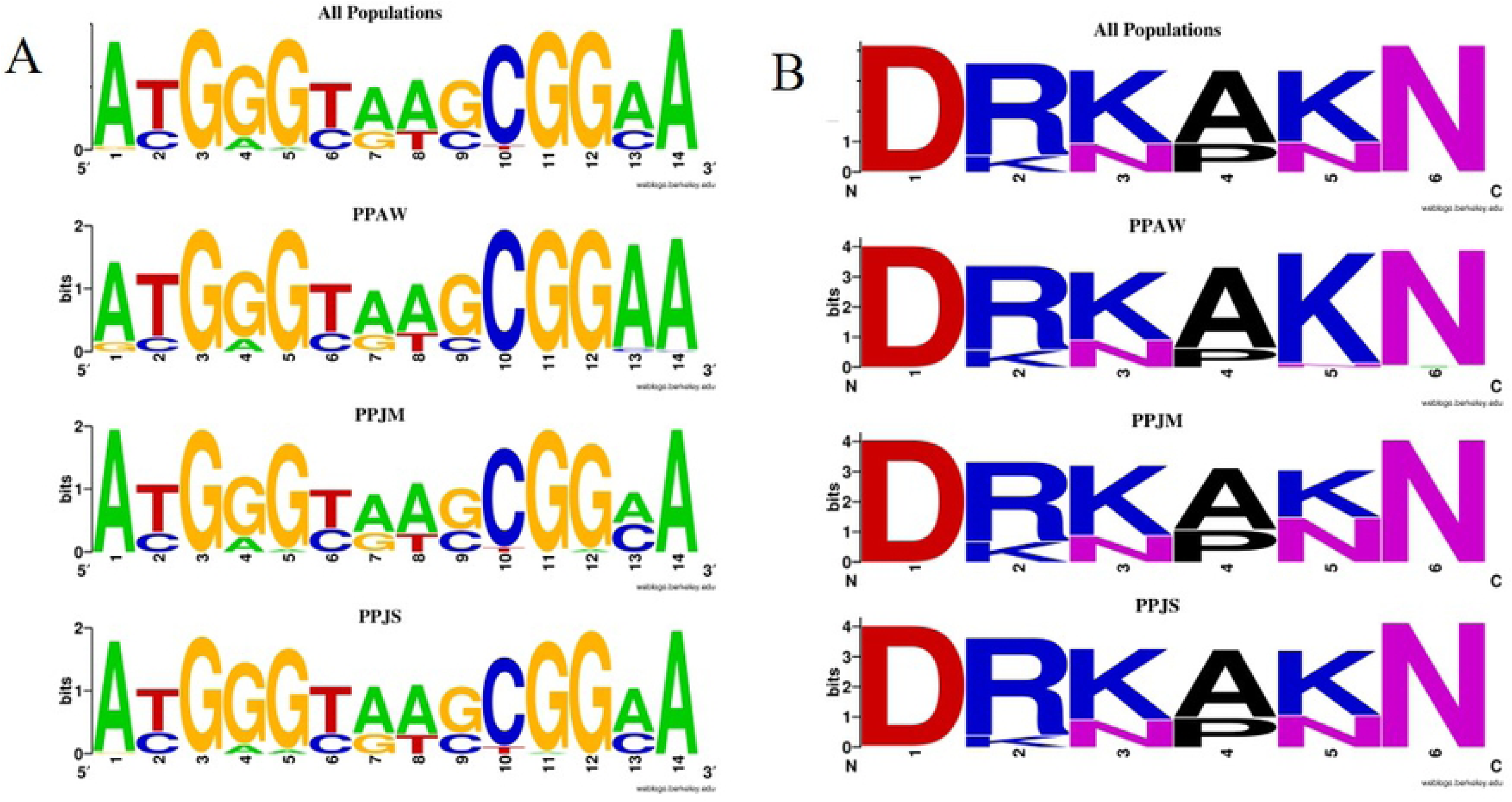
PpSP12 nucleotide and amino acid variation. (A) Weblogo illustrating the relative frequencies of nucleotide polymorphisms in wild caught *P. papatasi* populations from PPAW, PPJM, and PPJS. (B) Weblogo illustrating the relative frequencies of amino acid polymorphisms in wild caught *P. papatasi* populations from PPAW, PPJM, and PPJS.

#### Population genetics analysis

A total of 96 mature cDNA sequences were analyzed for *PpSP12* from Aswan (n=26), Malka (n=29), and Swaimeh (n=41). Twenty-nine haplotypes were identified with 14 variant sites. Of the 29 haplotypes, 22 were found in only one of the geographic study sites and 6 of the 7 shared haplotypes were found in 2 populations. One haplotype, H_1, was present in all 3 populations and was the most common haplotype. The Aswan, Egypt, population had 5 unique haplotypes (H_2, H_3, H_5, H_8, H_9) with 3 of those being private haplotypes (H_5, H_8, H_9). The Malka, Jordan, population exhibited 9 unique haplotypes (H_10, H_13, H_14, H_15, H_16, H_18, H_19, H_20, H_21) with 6 of the 9 being private haplotypes (H_10, H_14, H_15, H_18, H_19, H_20). The Swaimeh, Jordan, population demonstrated 8 unique haplotypes (H_22 to H_29) with 3 private haplotypes (H_22, H_24, H_25). A variety of population genetics parameters were assessed (Table 2) indicating genetic homogeneity for *PpSP12* across the three populations. Tajima’s D and Ka/Ks analysis indicated that this protein is not undergoing positive selection but rather it is either neutral or possibly experiencing purifying selection (Table 2). Furthermore, population structure is not indicated as Fst uncovered little genetic variability in pairwise comparisons (Table 3). The *PpSP12* median-joining network does not demonstrate any notable clustering separating the different populations from one another (Fig 3). Although there are 29 total haplotypes, the haplotypes are differentiated from one another by only one mutation. The Ka/Ks ratio, a diversifying selection index, was 0.293 or less across the sliding window analysis of the protein for all populations indicating purifying or stabilizing selection of this protein (Table 4).

**Fig 3.**
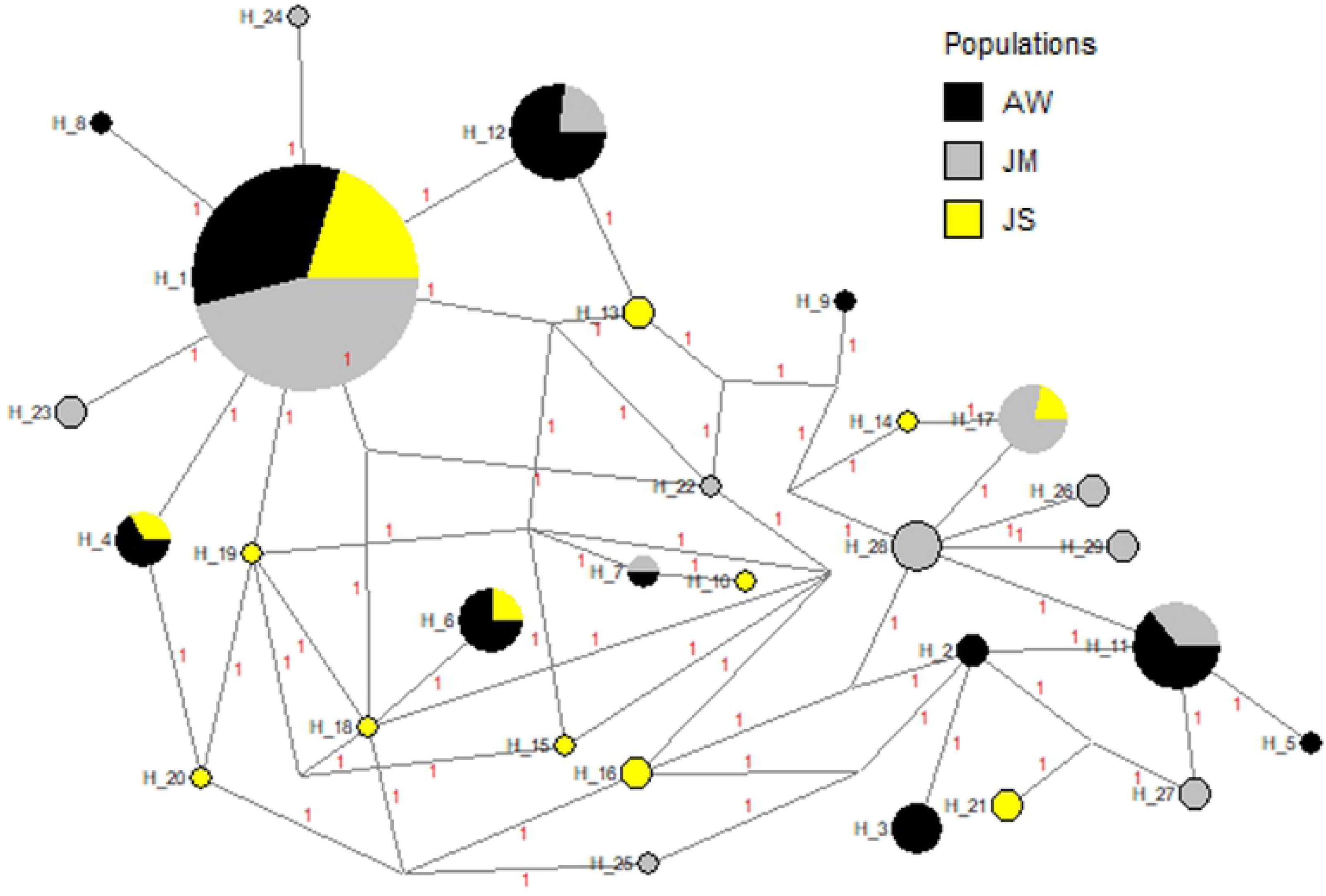
Median-joining network for PpSP12 *P. papatasi* haplotypes. Circle size and circle color indicates frequency and geographical location of haplotypes, respectively. Haplotype numbers are written next to the corresponding circle H_XX. Red numbers between haplotypes indicate number of mutations between haplotypes.

**Table 2.**
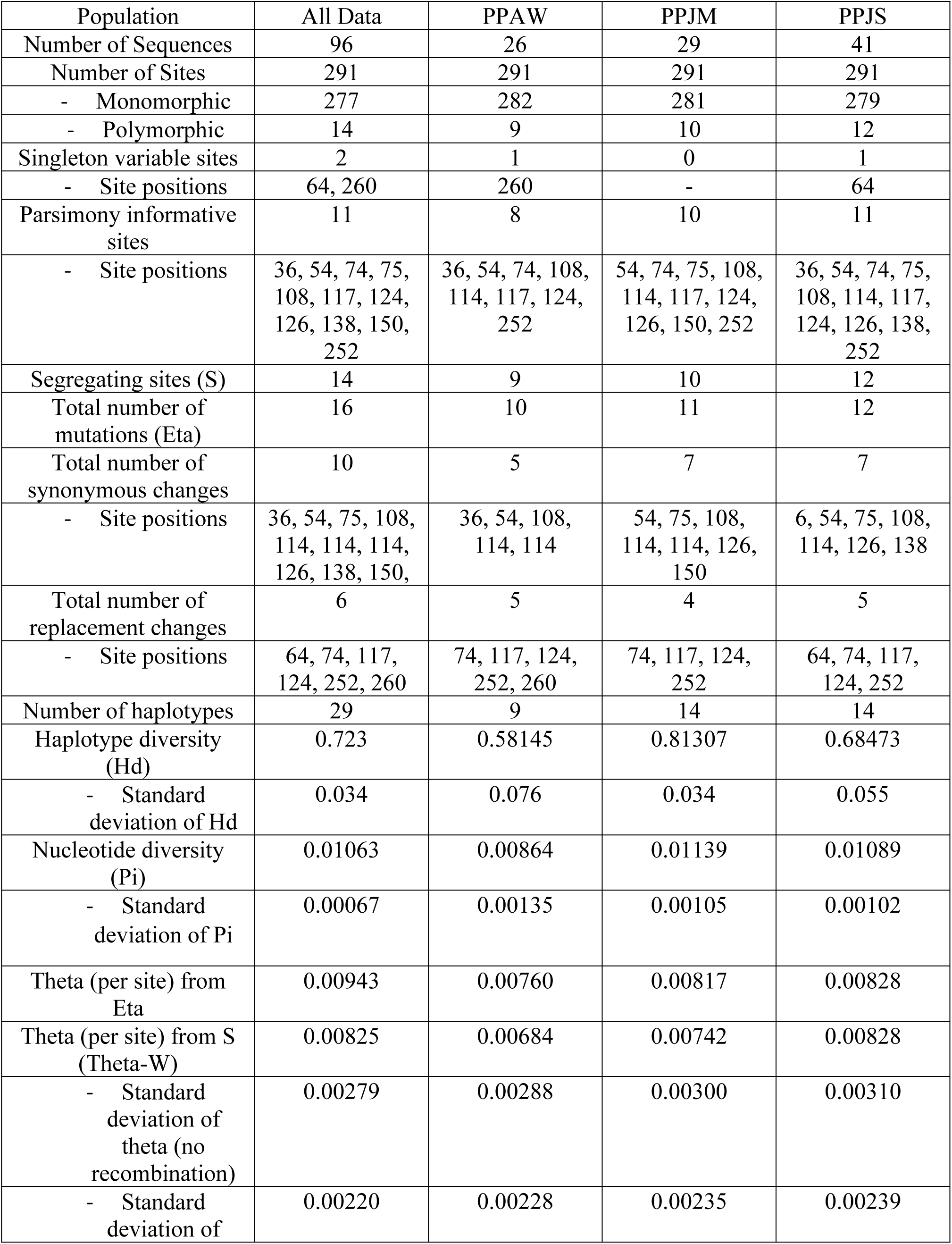
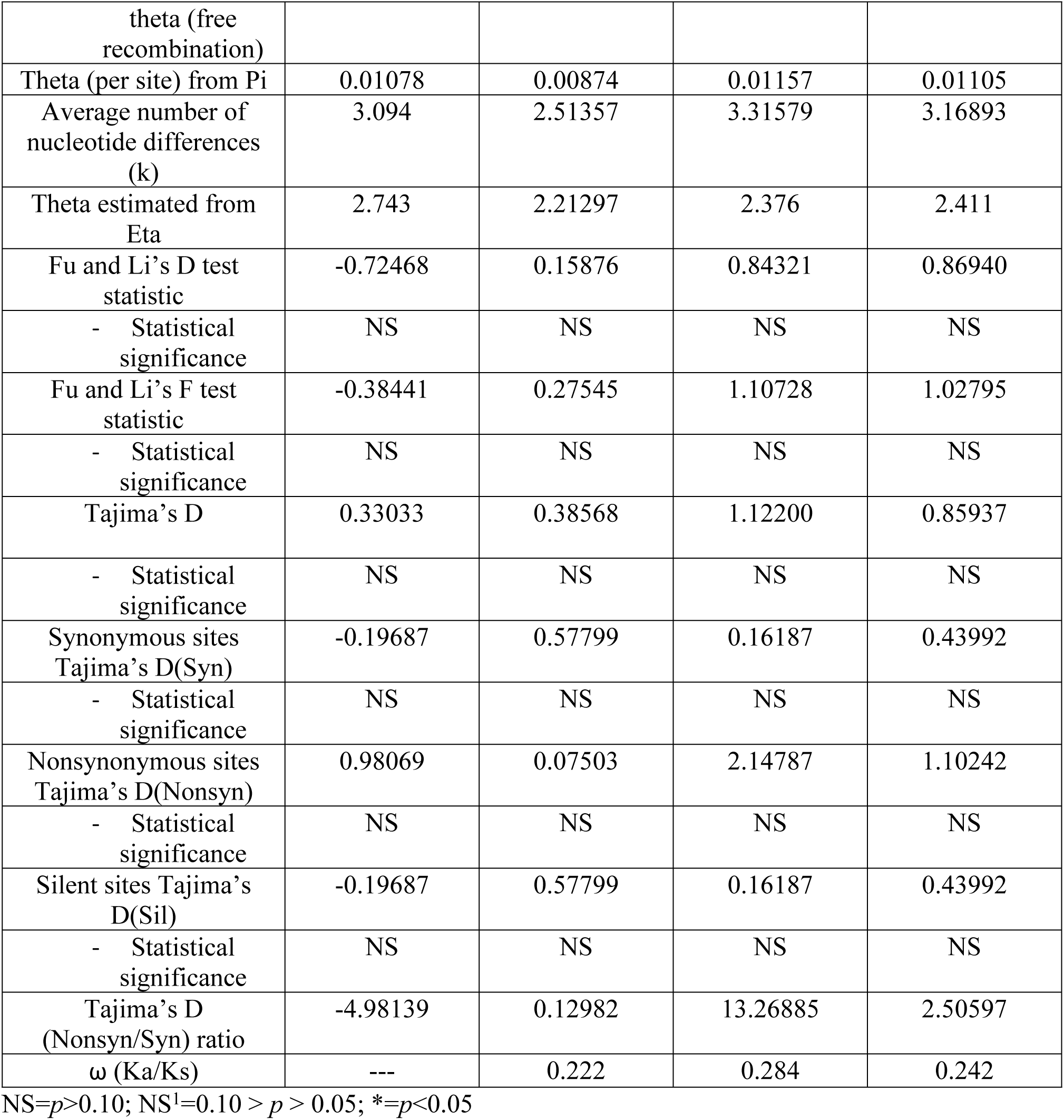
**PpSP12 population genetics analyses for *P. papatasi* populations**

**Table 3.** ***PpSP12* pairwise comparisons of genetic differentiation estimates.**

| POP 1 | POP 2 | Hs | Ks | Gst | Fst | Dxy | Da |
| --- | --- | --- | --- | --- | --- | --- | --- |
| PPAW | PPJM | 0.70381 | 2.93656 | 0.05362 | 0.06720 | 0.01074 | 0.00072 |
| PPAW | PPJS | 0.64501 | 2.91461 | 0.01242 | 0.03412 | 0.01011 | 0.00034 |
| PPJM | PPJS | 0.73758 | 3.22977 | 0.02492 | -0.00011 | 0.01114 | 0.00000 |

**Table 4.**
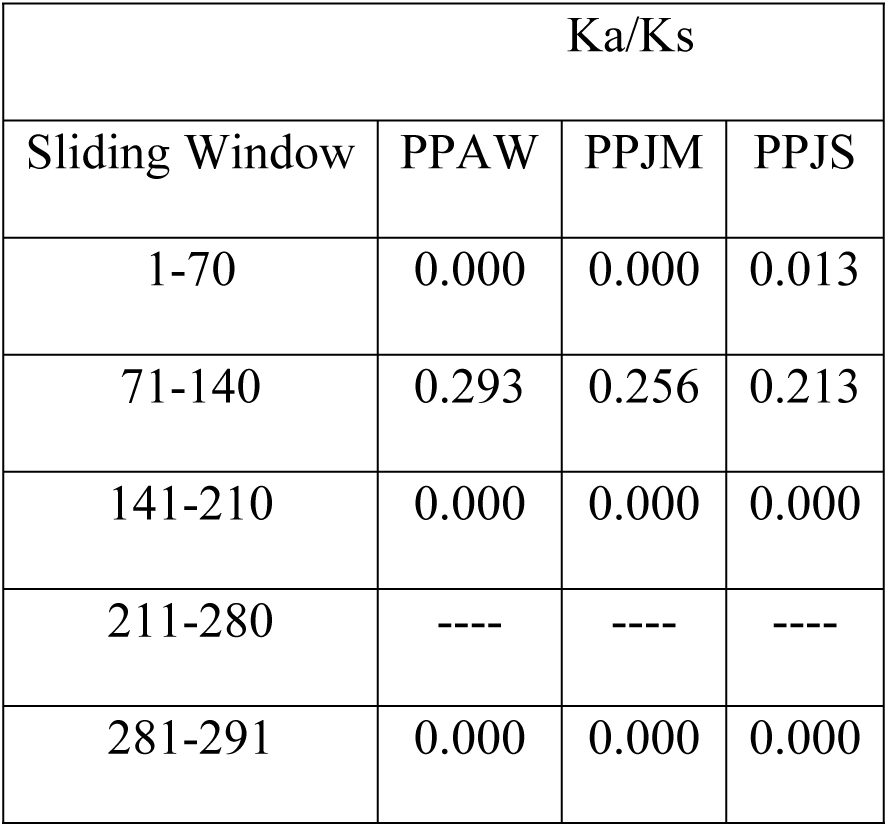
PpSP12 sliding window analysis. Ka/Ks were plotted for every 70 codons. Values greater than one suggest the potential for positive selection. ----indicates a lack of polymorphic data in the window to calculate a Ka/Ks value.

#### Secondary structure & T-cell epitope predictions

The mature amino acid sequence for PpSP12 predicted that only one polymorphic site (R60) was found in an α-helix whereas the other six polymorphic sites were found in predicted coils (D57, K74, A77, K119, N122) (Fig 4). All 6 polymorphic sites are found in predicted MHC II T-cell epitope binding sites though this should not interfere with the potential for T-cell activation as the polymorphic sites are found in the middle of the predicted binding sites and surrounded by conserved regions. Of the 140 amino acids included in this analysis, 95 amino acids were predicted to be potential epitope recognition sites. The areas of the amino acid sequence with the highest predicted binding affinities occur between the lysine residue at position 2 (K2) and the proline residue at position 23 (P23) as well as the tyrosine residue at position 110 (Y110) and the asparagine residue at position 138 (N138). Of the 78 total HLA alleles tested using the two software tools, all 51 alleles from ProPred identified potential binding sites though certain alleles, such as DRB1_03, DRB1_11, and DRB1_13, had greater binding affinities than the others. The alleles with the strongest binding affinity potential identified by IEDB software included DQA1_0401/DQB1_0402, DPA1_0103/DPB1_0201, and DRB1_0301. The DQA1/DQB1 and DPA1/DPB1 alleles demonstrated a greater affinity for residues between K2 to A20 and DRB1_0301 demonstrated a greater affinity for Y110 to F126, bookending the mature PpSP12.

**Fig 4.**
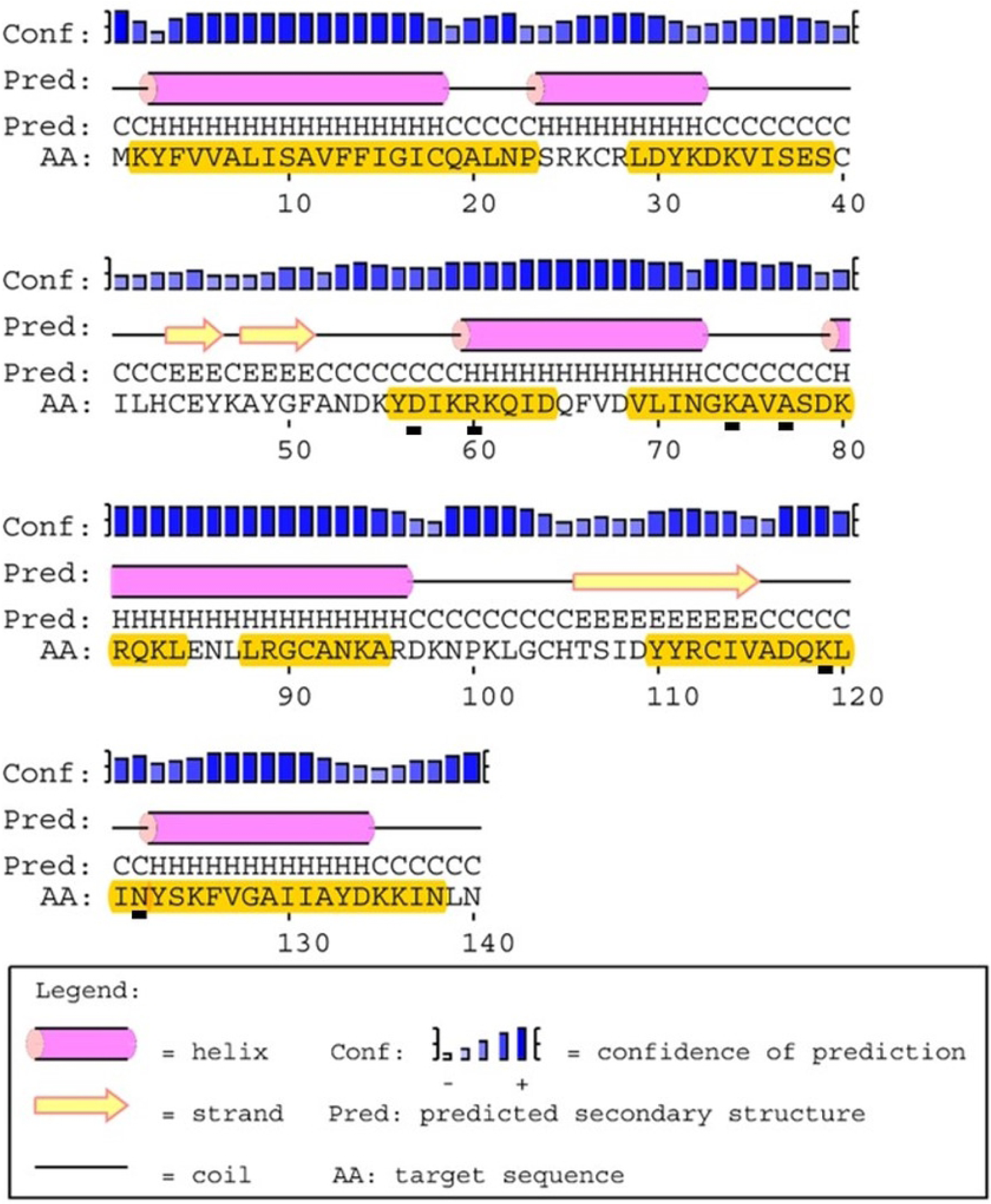
PpSp12 secondary structure, polymorphic sites, and MHC class II epitope predictions. The mature PpSP12 amino acid sequence predicted secondary structure. Yellow highlighted amino acids indicate the predicted MHC class II predicted promiscuous peptides. Individual amino acids underlined in black indicate unique polymorphic sites. Predicted secondary structure based on sequence accession #AGE83083[13].

### PpSP14 in-depth analyses

#### Nucleotide and amino acid genetic diversity

The 246-bp *PpSP14* fragment produced 23 polymorphic sites (Fig 5A). Similar to *PpSP12*, there exists limited heterogeneity between the populations studied. Position 11 demonstrates variation in all 3 populations with equal representation of adenine and guanine. In position 17, the Jordan populations have roughly equal rates of cytosine and guanine but the Egypt population has cytosine in the majority. Guanine dominates at position 21 in both Jordan populations but is equally represented with cytosine in the Egypt population. The remaining 20 polymorphic sites present similar levels of heterogeneity across all 3 populations. The translated PpSP14 amino acid sequence has 14 variable positions out of 82 total amino acids (Fig 5B). At position 1 and 3, leucine, valine, and isoleucine are all easily substituted for one another since they are hydrophobic and prefer to be buried in the protein core. In position 5, the Aswan, Egypt, population demonstrates limited substitution of the asparagine amino acid with lysine, both polar amino acids. Lysine and arginine are relatively equal for all populations at position 8. Lysine and arginine belong to the same basic amino acid group and are known substitutions for one another [60]. Serine and threonine, position 13, are also easily substituted for one another but more threonine is found in the Egypt population compared to the Jordan populations. At position 10, threonine and alanine substitutions are found in all populations but are more frequent in the Egypt population. Even though threonine is polar and alanine is nonpolar, both are small amino acids and threonine’s versatility of being inside or outside of the protein, the substitution can be functionally sound. At position 11, threonine is substituted by isoleucine in both Jordanian populations. Even though isoleucine is hydrophobic and threonine is polar, threonine may be on the inside of this protein making this substitution possible, similar to the substitution at position 10 [60].

**Fig 5.**
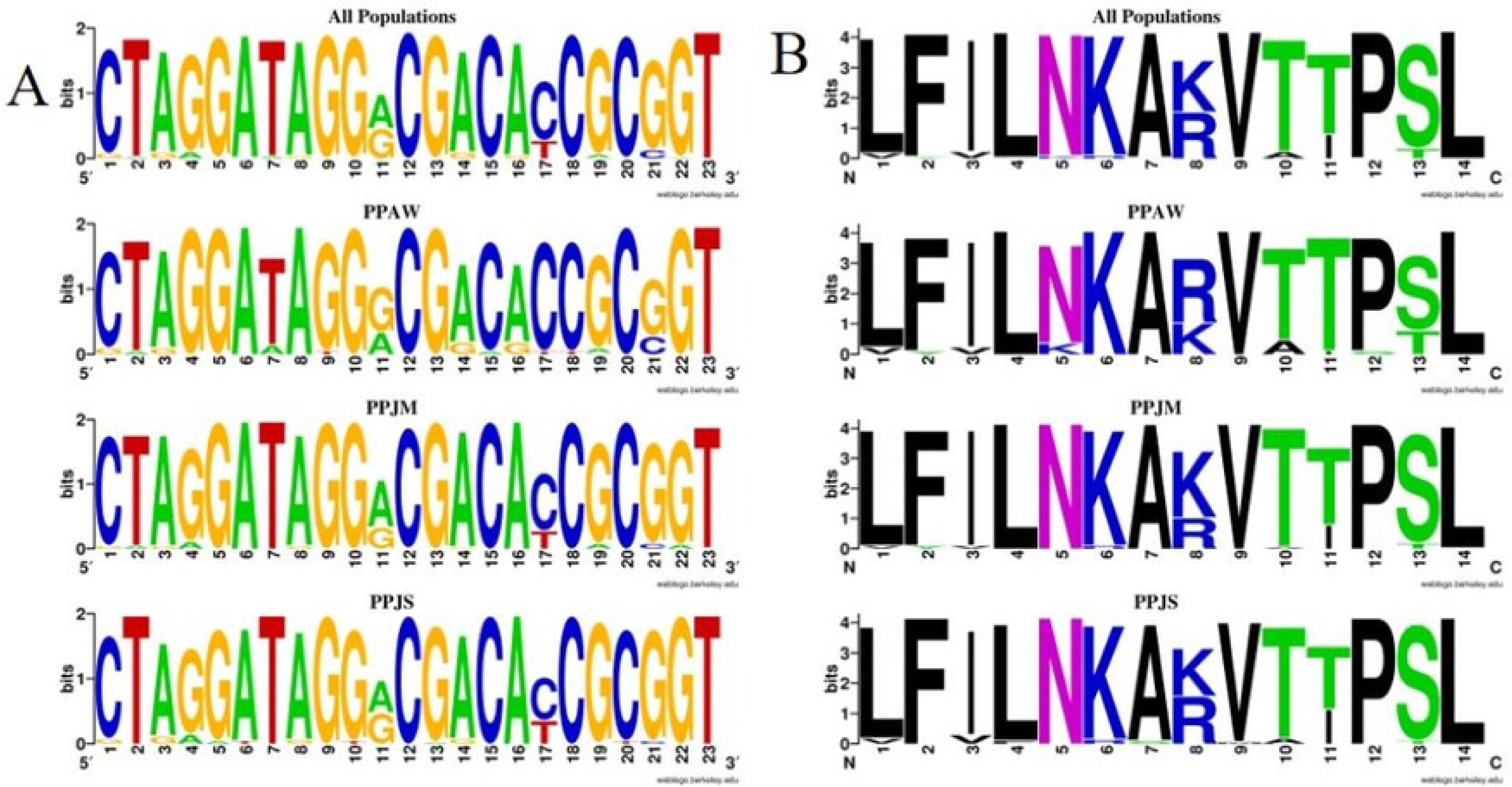
PpSP14 nucleotide and amino acid variation. (A) Weblogo illustrating the relative frequencies of nucleotide polymorphisms in wild caught *P. papatasi* populations from PPAW, PPJM, and PPJS. (B) Weblogo illustrating the relative frequencies of amino acid polymorphisms in wild caught *P. papatasi* populations from PPAW, PPJM, and PPJS.

#### Population genetics analysis

A total of 119 mature cDNA sequences were analyzed for *PpSP14* from PPAW (n=29), PPJM (n=44), and PPJS (n=46). Thirty-eight haplotypes 23 variant sites. Of the 38 haplotypes, 25 were found in only one of the geographic study sites, 8 were shared between PPAW/PPJM or PPJM/PPJS, but none were shared by PPAW/PPJS. Five haplotypes were present in all three populations (H_1, H_5, H_8, H_10, H_13), with H_5 the most common haplotype. PPAW had 6 unique haplotypes (H_3, H_7, H_9, H_11, H_12, H_14) with 1 of those designated a private haplotype (H_3). PPJM had 8 unique haplotypes (H_16, H_18, H_21, H_22, H_23, H_25, H_26, H_27) with 6 of the 8 being private haplotypes (H_16, H_18, H_22, H_23, H_25, H_26). PPJS had 11 unique haplotypes (H_28, H_29, H_30, H_31, H_32, H_33, H_34, H_35, H_36, H_37, H_38) with 6 of the 11 being private haplotypes (H_31, H_32, H_33, H_36, H_37, H_38). The population genetics assessment indicates genetic homogeneity for *PpSP14* across the 3 populations (Table 5). Although the Tajima’s D values were negative across all populations, the values were not significant and do not deviate far from zero indicating no selection. The majority of Ka/Ks values are under 1 or do not deviate far from 1 further indicating no selection acting on *PpSP14*. There is the potential for population structuring as Fst demonstrated moderate genetic differentiation between PPAW and PPJM (0.10771) and PPAW and PPJS (0.09091) and little genetic differentiation between PPJM and PPJS (0.02346) (Table 6). The *PpSP14* median joining network does not demonstrate any significant clustering separating the different populations from one another (Fig 6). The PPJS haplotypes might be clustering together as compared to PPAW and PPJM but the 38 haplotypes are differentiated from one another by only one mutation. The only exception being haplotypes H_7 and H_12, both from PPAW are differentiated by 3 mutations from one another. The Ka/Ks ratio in the sliding window from 141-210 indicated potential positive selection in both PPJM and PPJS populations but not in the PPAW population (Table 7).

**Table 5.**
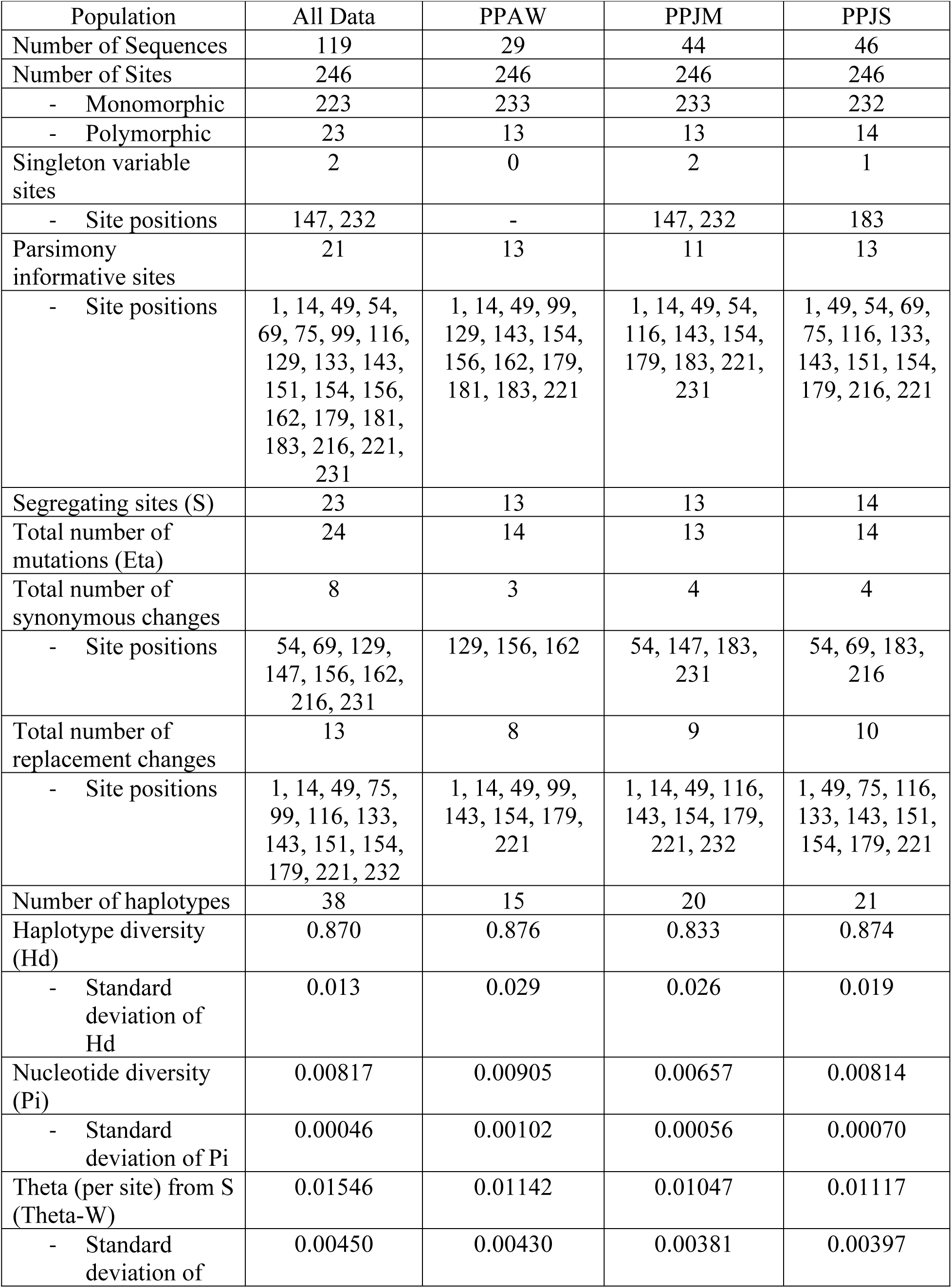
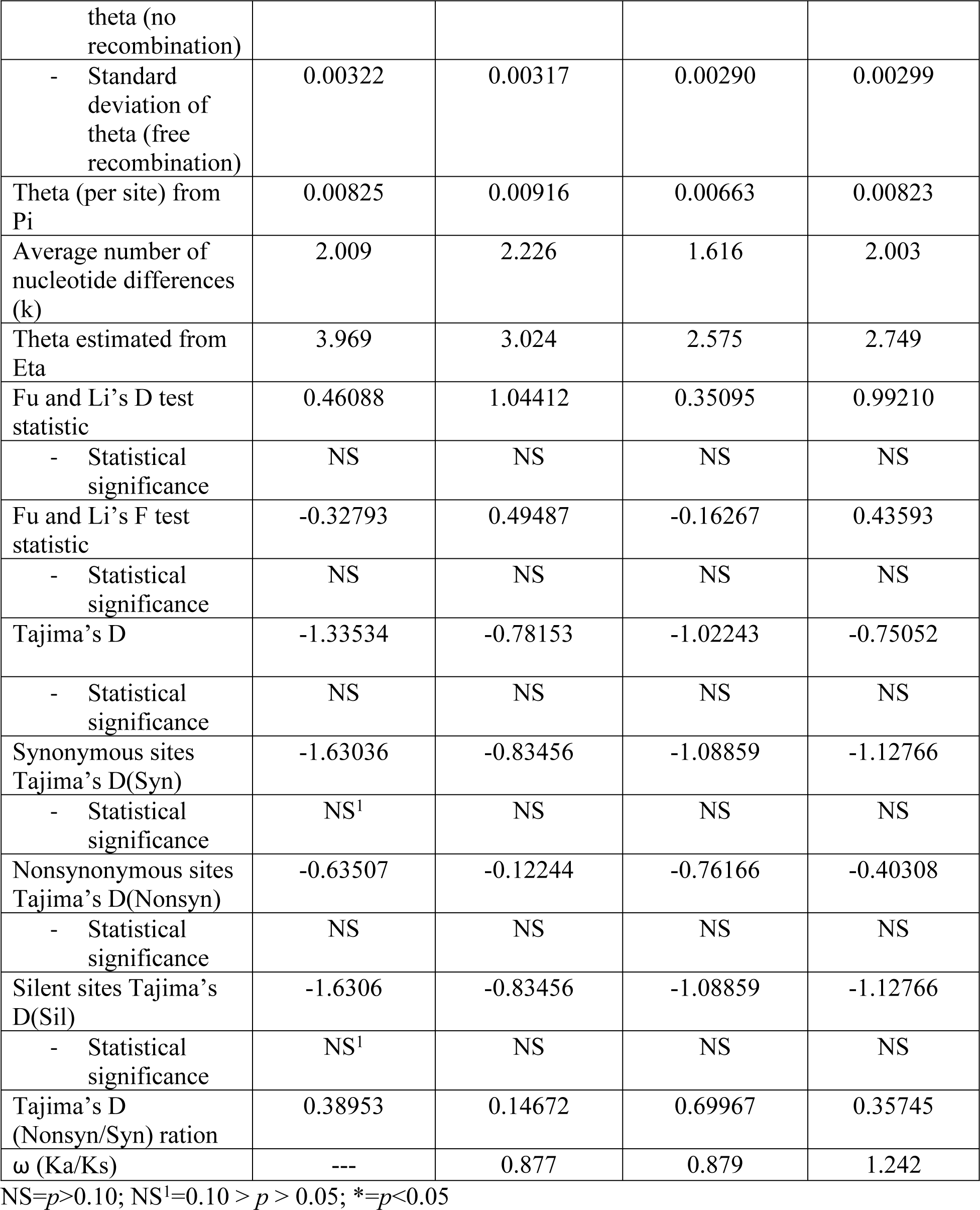
**PpSP14 population genetics analyses for *P. papatasi* populations.**

**Fig 6.**
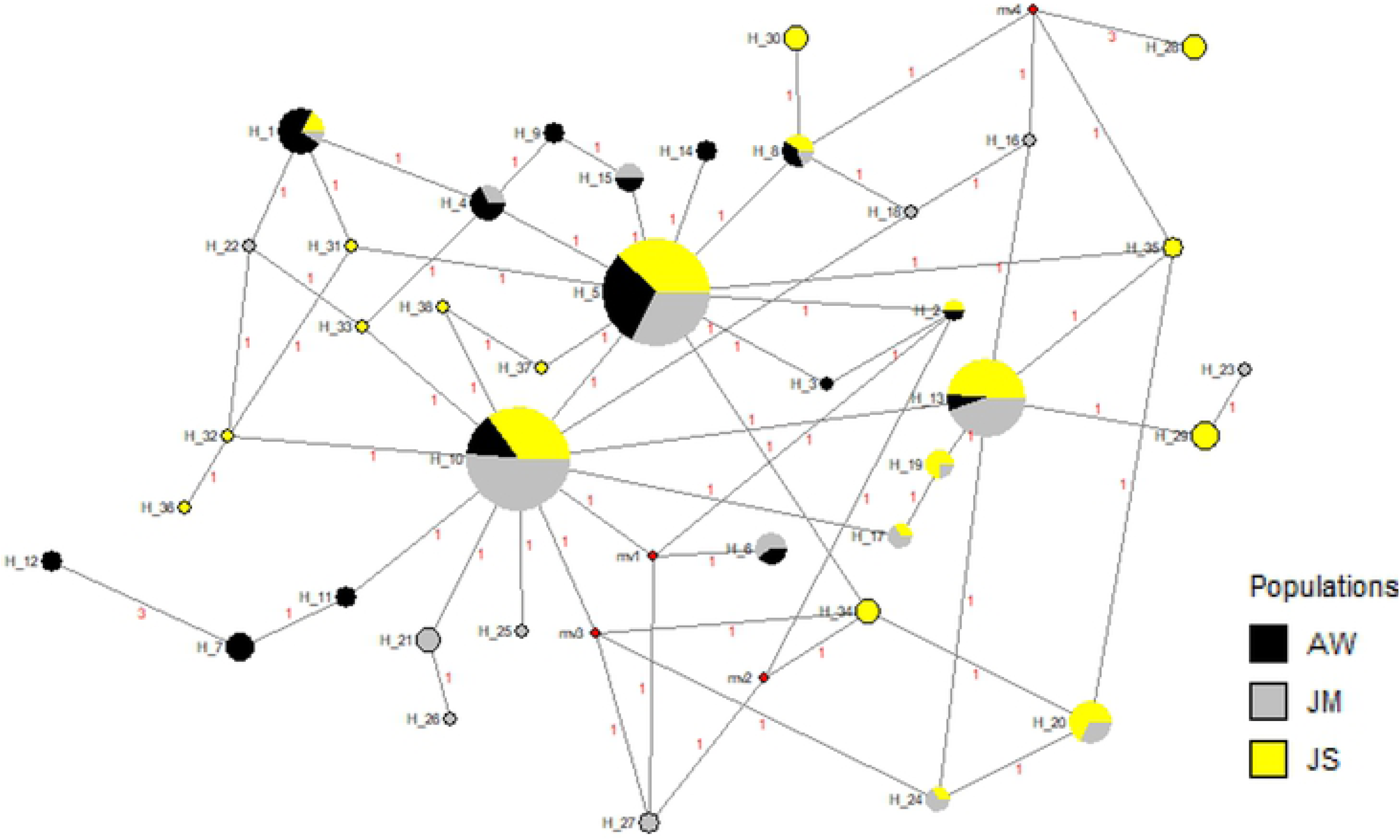
Median-joining network for PpSP14 *P. papatasi* haplotypes. Circle size and circle color indicates frequency and geographical location of haplotypes, respectively. Haplotype numbers are written next to the corresponding circle H_XX. Red numbers between haplotypes indicate number of mutations between haplotypes.

**Table 6.**
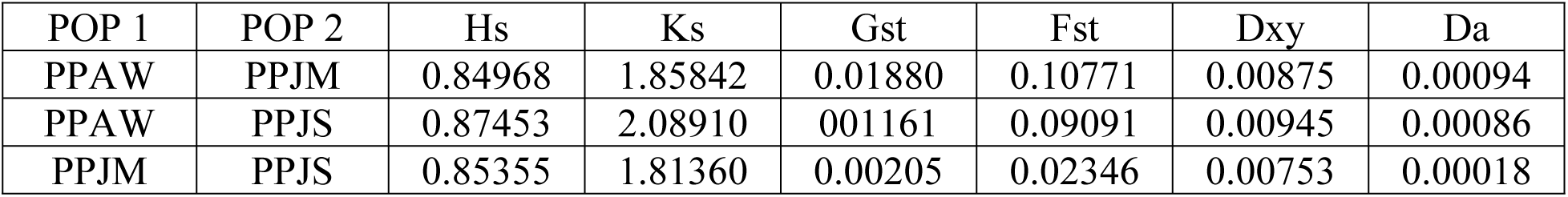
**PpSP14 pairwise comparisons of genetic differentiation estimates.**

**Table 7.**
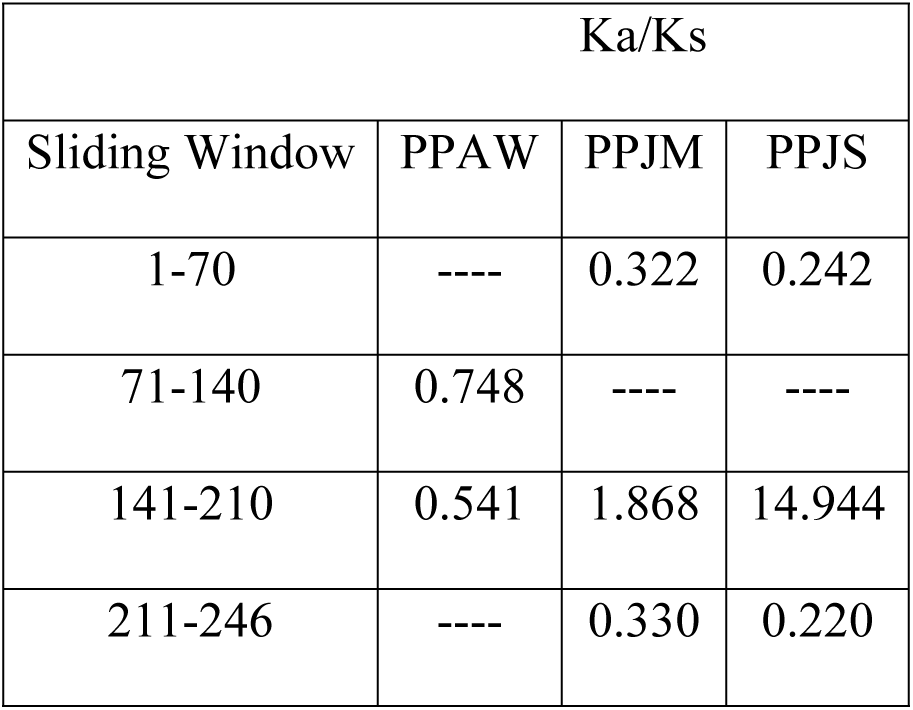
**PpSP14 sliding window analysis.** Ka/Ks were plotted for every 70 codons. Values greater than one suggest the potential for positive selection. ----indicates a lack of polymorphic data in the window to calculate a Ka/Ks value.

| Ka/Ks |  |  |  |
| --- | --- | --- | --- |
| Sliding Window | PPAW | PPJM | PPJS |
| 1-70 | ---- | 0.322 | 0.242 |
| 71-140 | 0.748 | ---- | ---- |
| 141-210 | 0.541 | 1.868 | 14.944 |
| 211-246 | ---- | 0.330 | 0.220 |

#### Secondary structure & T-cell epitope predictions

The mature amino acid sequence for PpSP14 predicted that 6 polymorphic sites (L65, K79, A85, K88, V91, T92) were found in α-helices whereas the other 8 polymorphic sites were found in predicted coils (L41, F45, I57, N73, T100, P101, S114, L118) (Fig 7). Twelve of the 14 polymorphic sites were found in predicted MHC II T-cell epitope binding sites. This variability may affect the predicted binding sites found between amino acids L41 and K58 and between amino acids L87 and C103, as the variable sites are found at the beginning and end of the fragment. There are no polymorphic sites found between amino acids M1 and F19. The variable sites found between H61 and A77 and I106 and T134 are found in the middle of the amino acid fragment and should not hinder binding. Of the 142 amino acids included in this analysis, 100 amino acids were predicted to be potential epitope recognition sites. The software prediction tools, IEDB and ProPred, agree that the areas of the amino acid sequence with the highest predicted binding affinities occur between methionine residue at position 1 (M1) and the phenylalanine residue at position 19 (F19) as well as the isoleucine residue at position 106 (I106) and the threonine residue at position 134 (T134). Similar to PpSP12, all 51 alleles from ProPred identified potential binding sites, particularly between residues M1 and F19. Alleles DRB1_04XX, DRB1_08XX, DRB1_11XX, DRB1_13XX, and DRB1_15XX had the highest binding affinities overall. To a lesser extent, the following alleles were also identified DRB1_010X, DRB1_030X, DRB1_070X, and DRB5_010X. The alleles with the strongest binding affinity potential identified by IEDB software included DRB3_0101, DPA1_0301/DPB1_0402, DRB1_1101, D_0101/DQB1_0501, DPA1_0103/DPB1_0201, DRB1_0301, DPA1_0201/DPB1_0101, and DRB5_0101.

**Fig 7.**
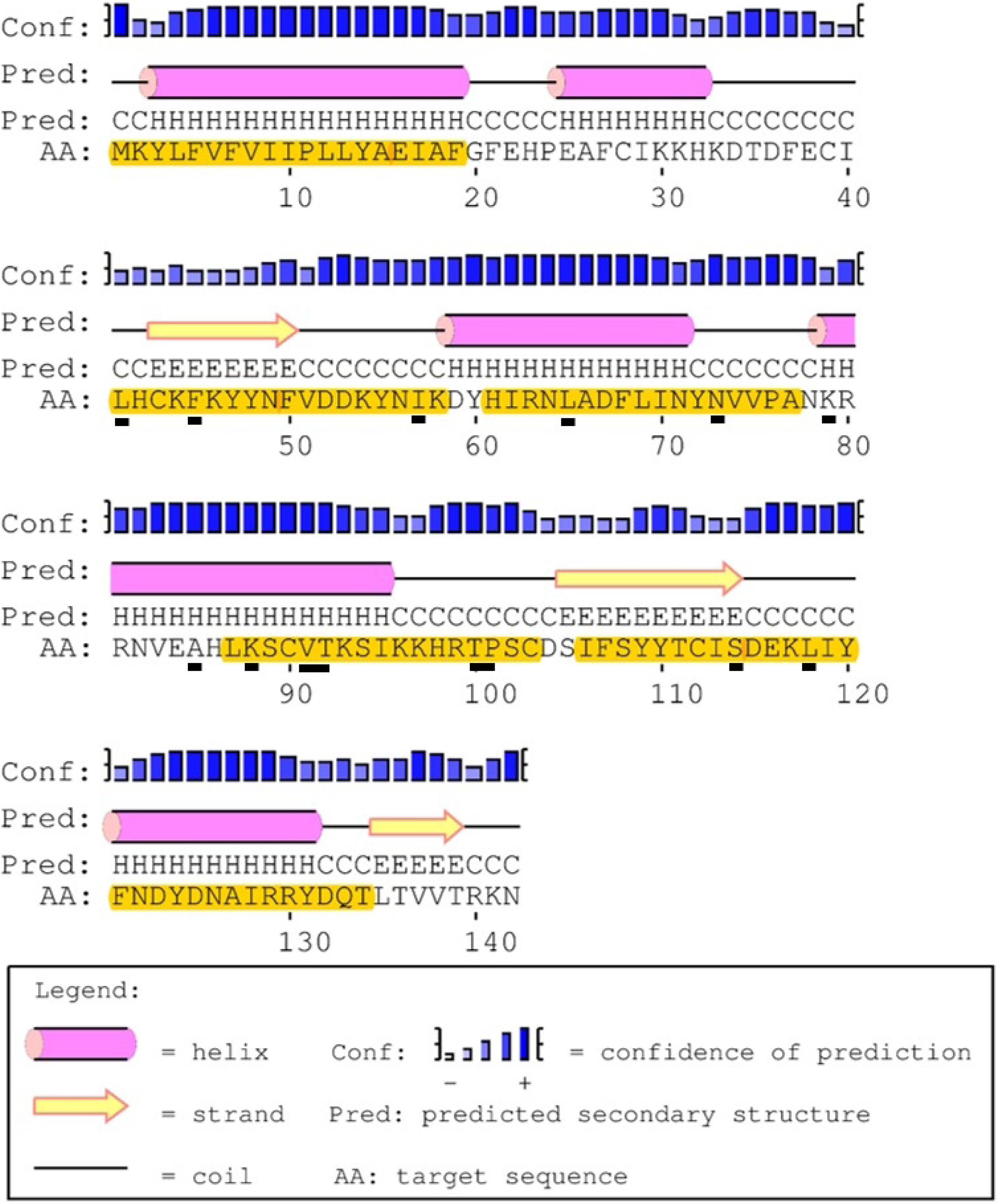
PpSp14 secondary structure, polymorphic sites, and MHC class II epitope predictions. The mature PpSP14 amino acid sequence predicted secondary structure. Yellow highlighted amino acids indicate the predicted MHC class II predicted promiscuous peptides. Individual amino acids underlined in black indicate unique polymorphic sites. Predicted secondary structure based on sequence accession #AGE83089[13].

### PpSP28 In-depth Analyses

#### Nucleotide and amino acid genetic diversity

The 651-bp *PpSP28* fragment produced 95 polymorphic sites (Fig 8A). Approximately, 61% of the polymorphic sites are transition substitutions and 26% are transversions. The remaining 12% of polymorphic sites are mostly conserved as the number of substitutions are so few. Positions 50 and 90 have three possible options at this site, each site a different combination of guanine, cytosine, adenine, or thymine. The translated PpSP28 amino acid sequence has 53 variable positions out of 184 total amino acids (Fig 8B). Eighteen of the variable sites demonstrate limited heterogeneity while the other 35 sites demonstrate significant variation between the populations and an abundance of amino acid substitutions. PpSP28 exhibits the greatest nucleotide and amino acid sequence variability of all 9 salivary proteins studied (Fig 8).

**Fig 8.**
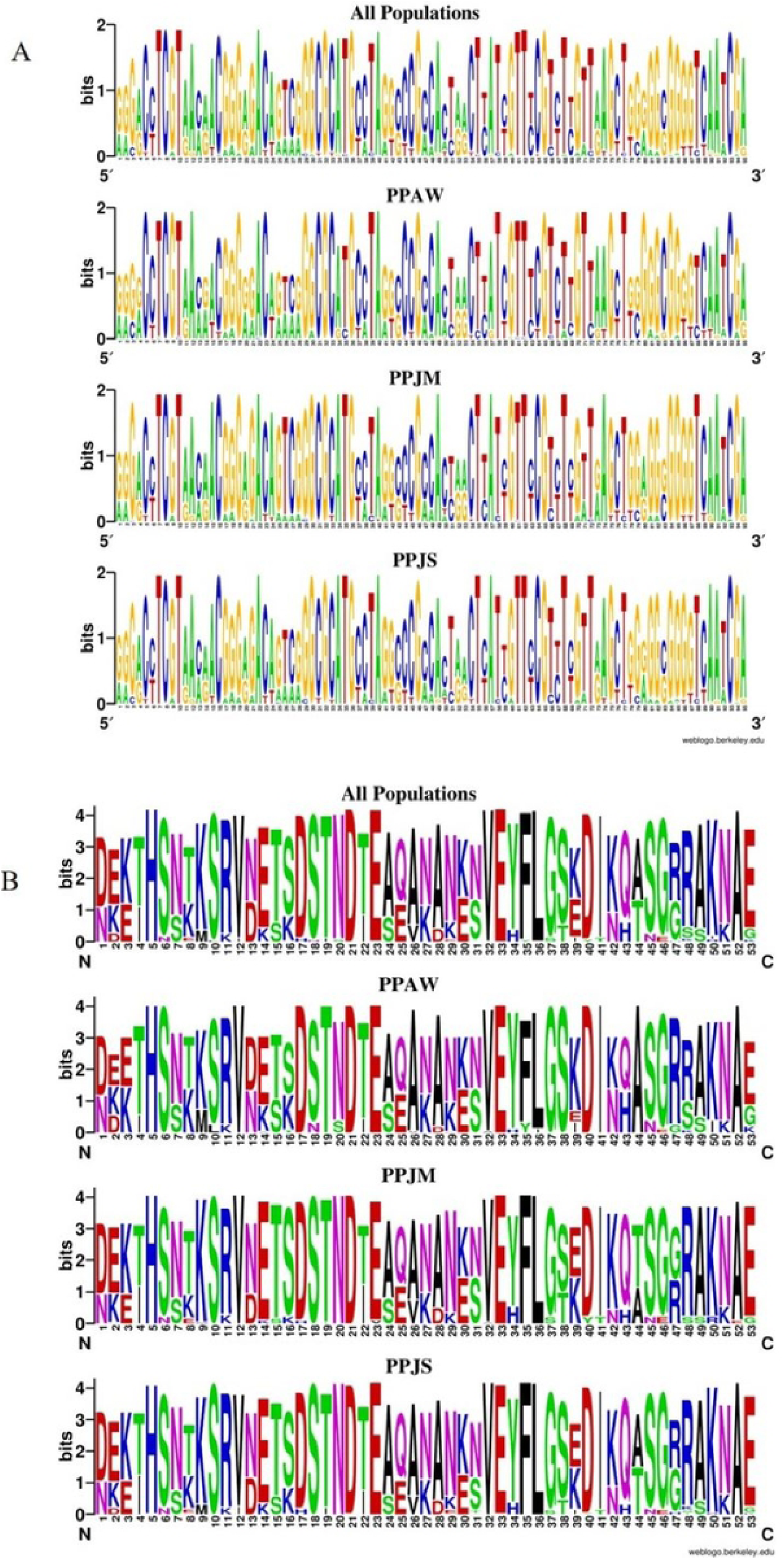
PpSP28 nucleotide and amino acid variation. (A) Weblogo illustrating the relative frequencies of nucleotide polymorphisms in wild caught *P. papatasi* populations from PPAW, PPJM, and PPJS. (B) Weblogo illustrating the relative frequencies of amino acid polymorphisms in wild caught *P. papatasi* populations from PPAW, PPJM, and PPJS.

#### Population genetics analysis

A total of 111 mature cDNA sequences were analyzed for *PpSP28* from Aswan (n=26), Malka (n=30), and Swaimeh (n=55). Ninety-five variant sites were identified in 122 haplotypes. Swaimeh, Jordan, tallied the most unique haplotypes (62) with 46 identified as private. Malka, Jordan, had 21 unique haplotypes with 16 identified as private. Aswan, Egypt, totaled 30 unique haplotypes with 24 identified as private. One haplotype was shared by all 3 populations (H_1) and 9 haplotypes were shared by 2 populations. As with *PpSP12* and *PpSP14*, various population genetics parameters were assessed (Table 8) indicating heterogeneity among the populations. Although significant variation is present in *PpSP28*, the analyses do not indicate that positive selection is acting on this salivary protein (Table 8). Population pairwise comparisons, like Fst, reveal great genetic differentiation, according to Wright (1978) between Aswan, Egypt, and Malka, Jordan at 0.10913, and moderate genetic differentiation between Aswan, Egypt, and Swaimeh, Jordan at 0.06936 (Table 9). There is little genetic differentiation between Malka, Jordan, and Swaimeh, Jordan, at 0.01595. The median joining network for *PpSP28* similarly does not exhibit any clear clustering of the Egypt or Jordan populations, but there are as many as 11 mutations separating connected haplotypes (Fig 9). *PpSP28* sliding window analysis of Ka/Ks demonstrates the potential for *PpSP28* to be under diversifying selection in several areas in contrast to the majority of the protein under purifying selection in all populations (Table 10). Values higher than one were detected in all 3 populations with PPJM having two sliding window regions with values over one compared to one sliding window region in PPJS and PPAW (Table 10).

**Table 8.**
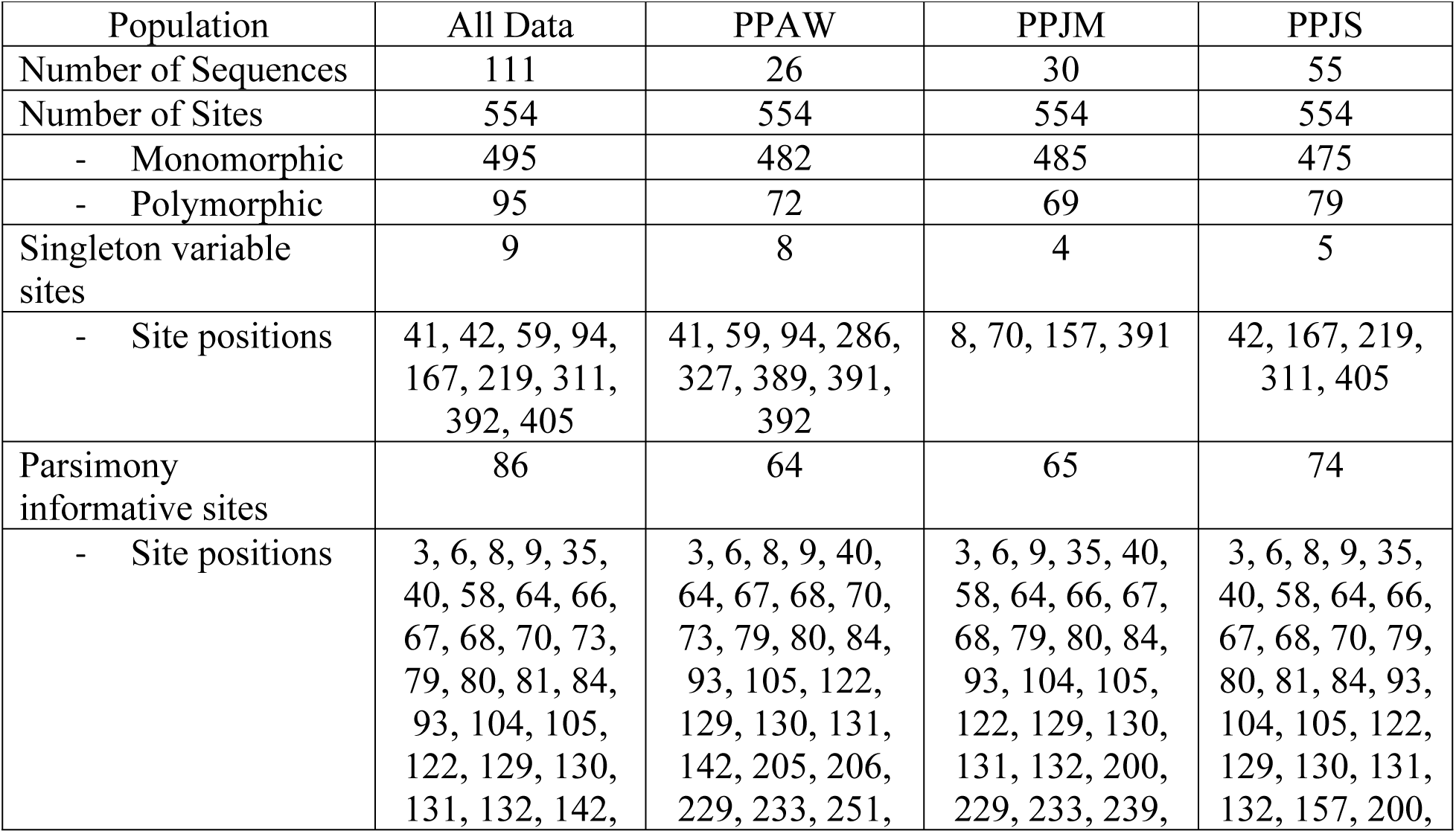
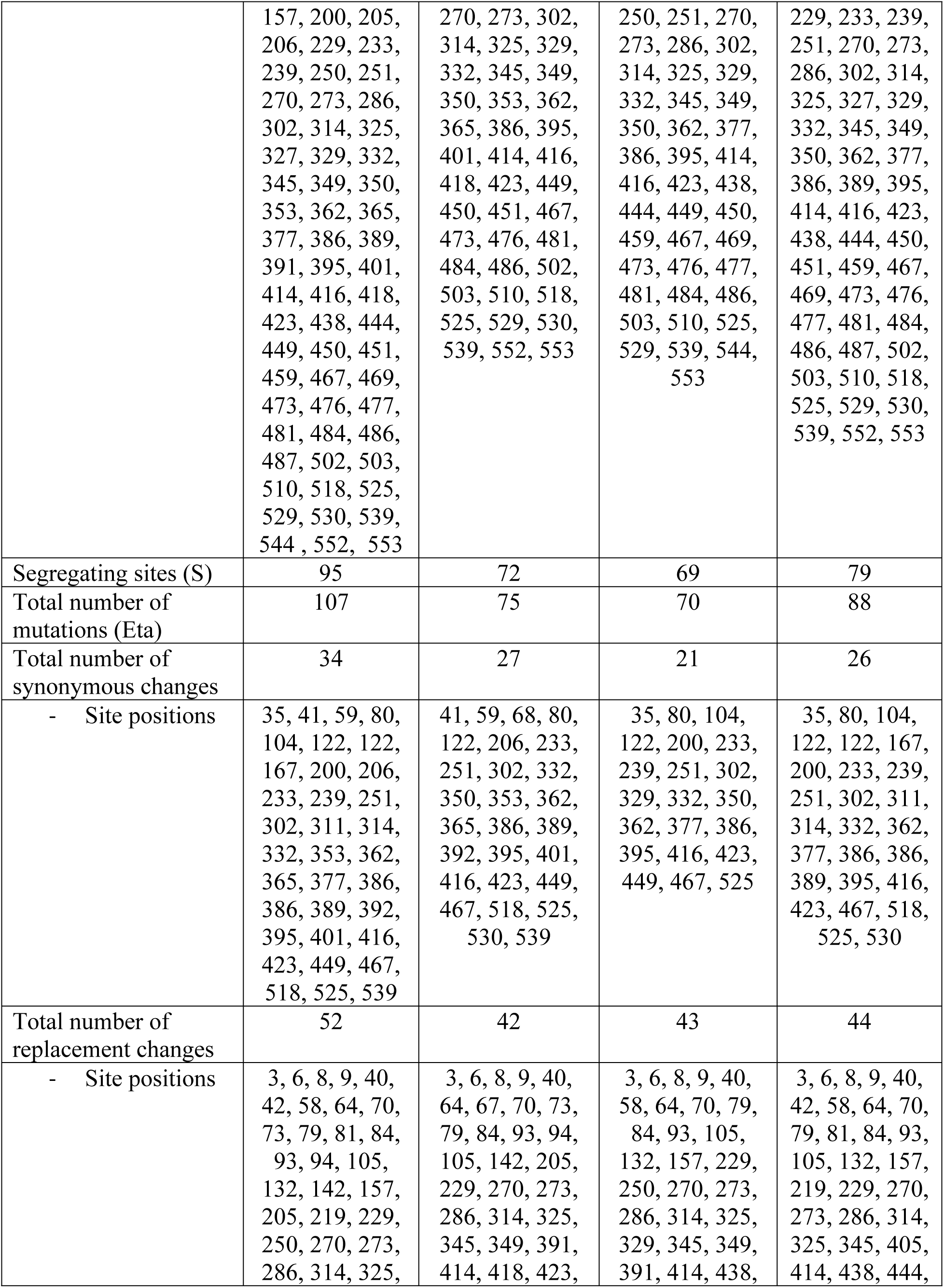
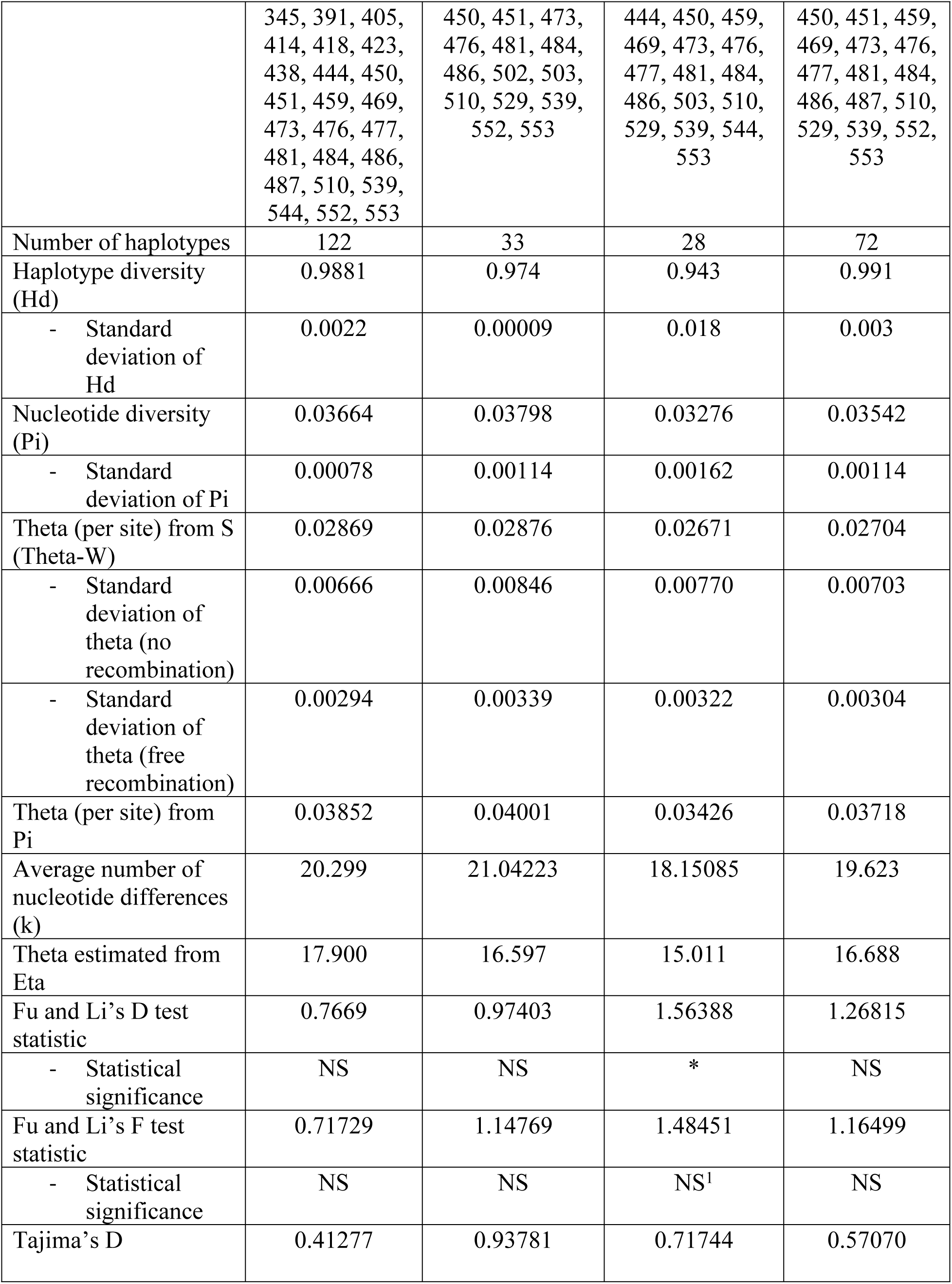
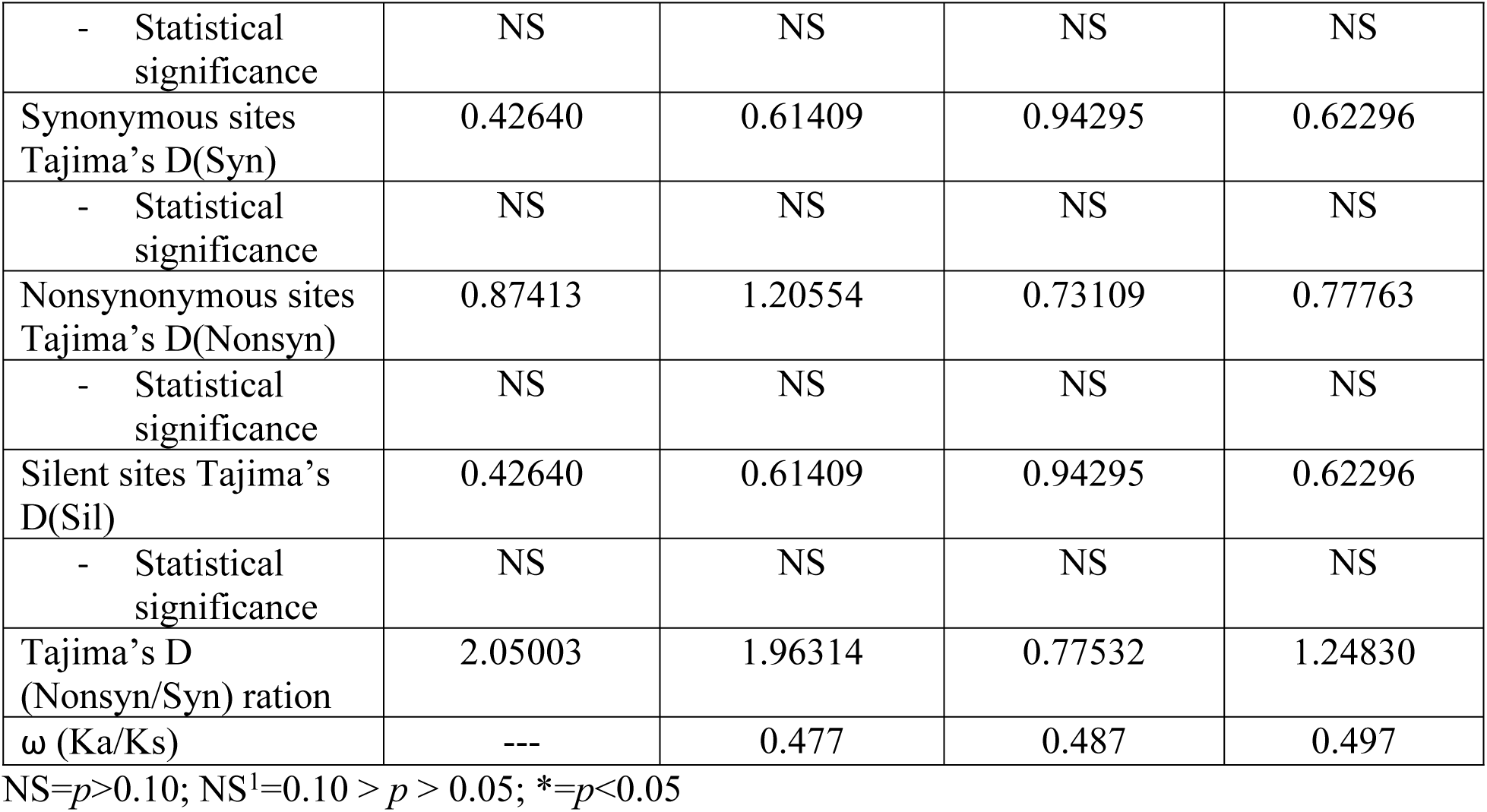
**PpSP28 population genetics analyses for *P. papatasi* populations**.

**Table 9.**
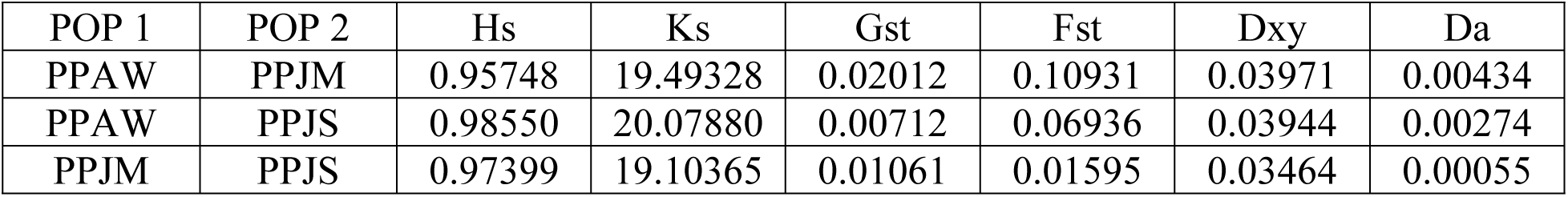
***PpSP28* pairwise comparisons of genetic differentiation estimates.**

| POP 1 | POP 2 | Hs | Ks | Gst | Fst | Dxy | Da |
| --- | --- | --- | --- | --- | --- | --- | --- |
| PPAW | PPJM | 0.95748 | 19.49328 | 0.02012 | 0.10931 | 0.03971 | 0.00434 |
| PPAW | PPJS | 0.98550 | 20.07880 | 0.00712 | 0.06936 | 0.03944 | 0.00274 |
| PPJM | PPJS | 0.97399 | 19.10365 | 0.01061 | 0.01595 | 0.03464 | 0.00055 |

**Fig 9.**
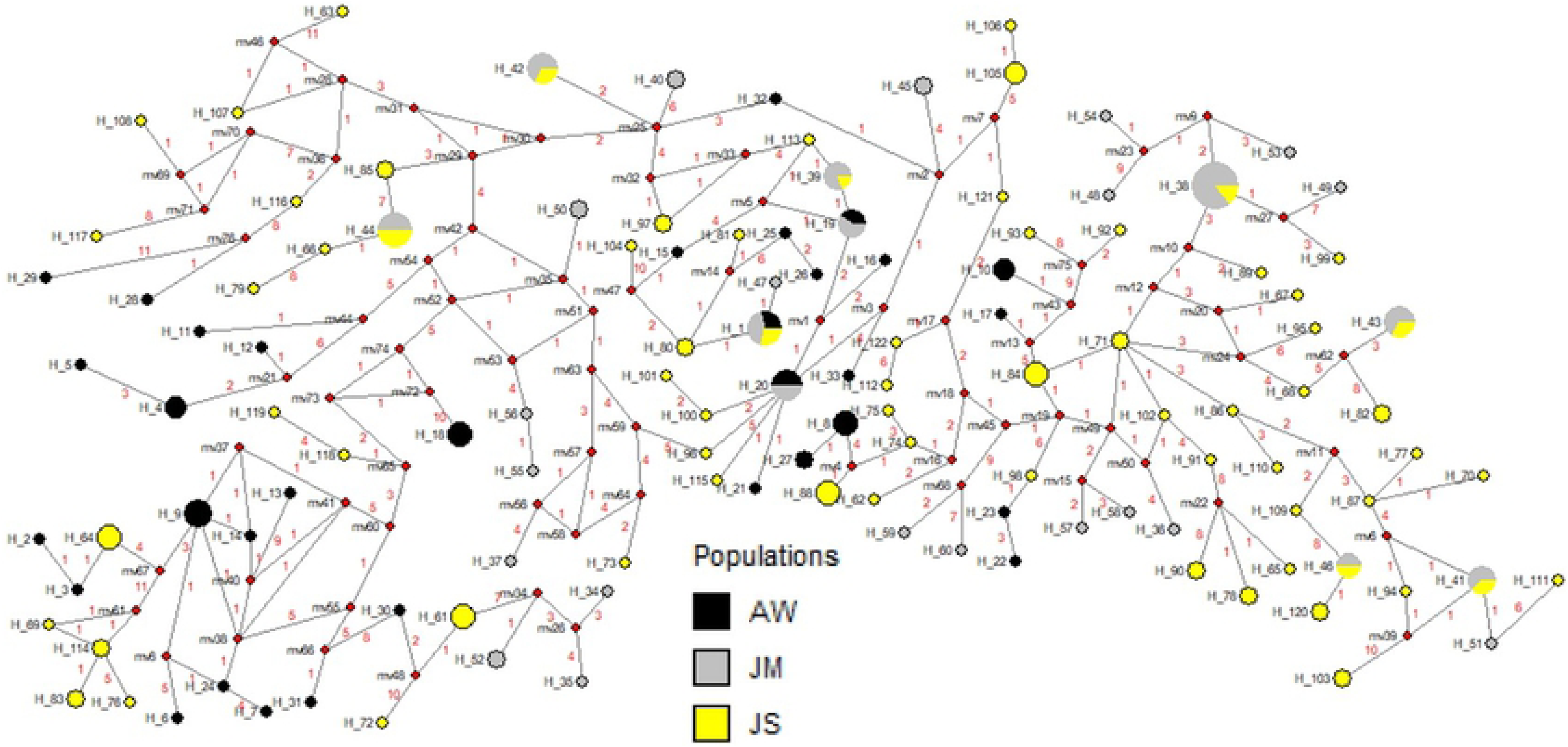
Median-joining network for PpSP28 *P. papatasi* haplotypes. Circle size and circle color indicates frequency and geographical location of haplotypes, respectively. Haplotype numbers are written next to the corresponding circle H_XX. Red numbers between haplotypes indicate number of mutations between haplotypes.

**Table 10.** PpSP28 sliding window analysis. Ka/Ks were plotted for every 70 codons. Values greater than one suggest the potential for positive selection.

| Ka/Ks |  |  |  |
| --- | --- | --- | --- |
| Sliding Window | PPAW | PPJM | PPJS |
| 1-71 | 1.506 | 1.329 | 1.437 |
| 72-141 | 0.624 | 0.427 | 0.482 |
| 142-211 | 0.413 | 0.059 | 0.254 |
| 212-281 | 0.453 | 0.473 | 0.571 |
| 282-351 | 0.852 | 0.570 | 0.756 |
| 352-421 | 0.031 | 0.038 | 0.024 |
| 422-491 | 0.391 | 0.890 | 0.862 |
| 492-554 | 0.845 | 1.333 | 0.534 |

#### Secondary structure & T-cell epitope predictions

The mature amino acid sequence for PpSP28 predicted a total of 53 polymorphic sites with 30 found in α-helices whereas the other 23 polymorphic sites were found in predicted coils (Fig 10). Twenty of the polymorphic sites are found in predicted MHC II T-cell epitope binding sites. Only 1 high affinity predicted binding site between residues M1 and S19 showed no variation. IEDB identified two alleles would recognize this region including DPA1_0301/DPB1_0402 and DPA1_01/DPB1_0401. The majority of the ProPred alleles recognized some combination of residues between M1 and Q28. Another conserved region was identified between residues F33 and S41 and was recognized by DRB1_08XX, DRB1_11XX, and DRB1_13XX. The final region demonstrating conservation between residues L46 and L54 was recognized but by very few alleles. Two of the regions with the strongest affinities housed the most variation such as residues L71 to Q89, with 9 variant sites, and F195 to F203, with 4 variant sites. The high degree of variation within potential epitope binding sites and the decreased variety of alleles identified should exclude PpSP28 from further vaccine development.

**Fig 10.**
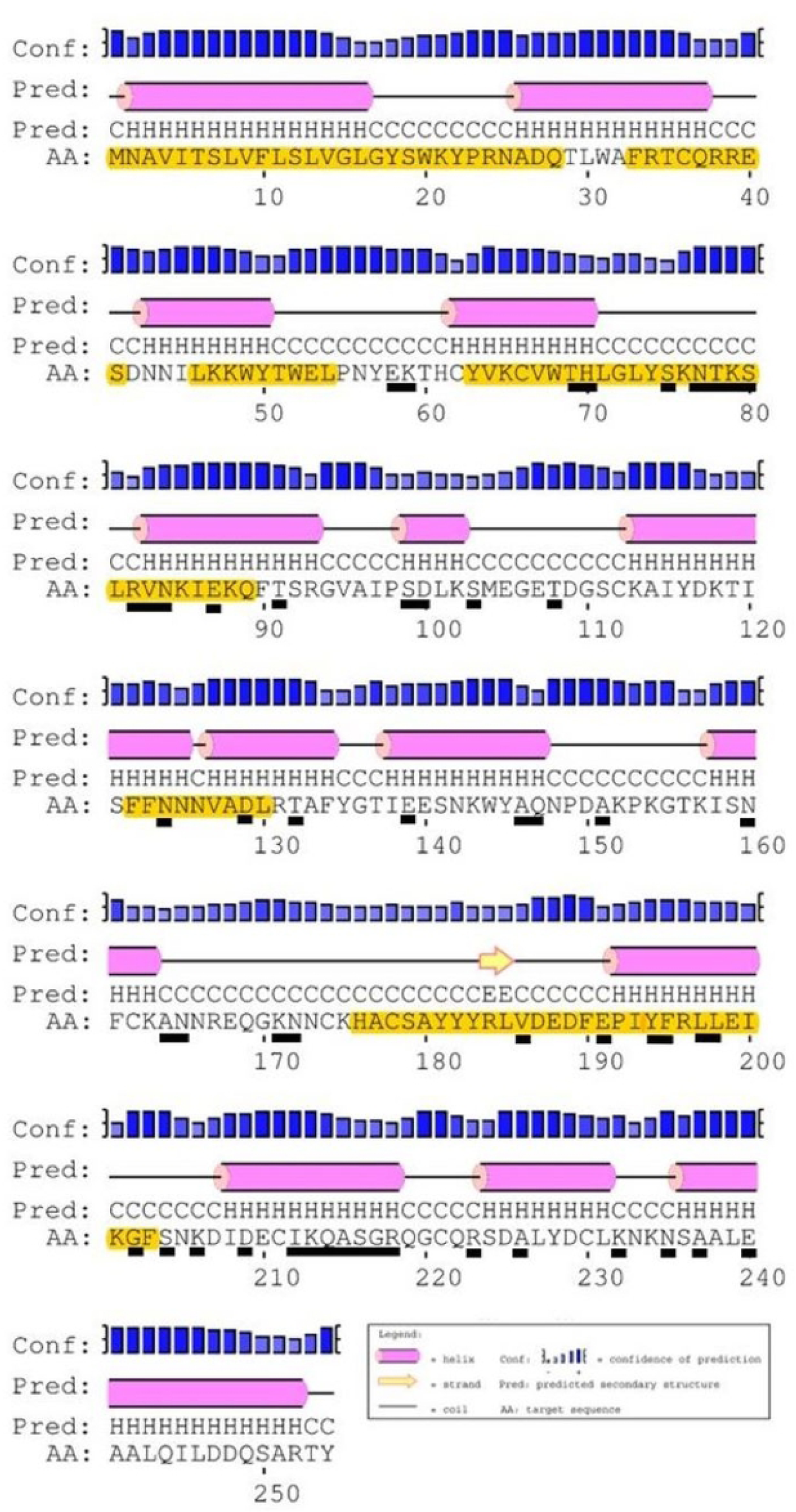
PpSp28 secondary structure, polymorphic sites, and MHC class II epitope predictions. The mature PpSP28 amino acid sequence predicted secondary structure. Yellow highlighted amino acids indicate the predicted MHC class II predicted promiscuous peptides. Individual amino acids underlined in black indicate unique polymorphic sites. Predicted secondary structure based on sequence accession #AGE83090 and #AGE83091[13].

### Multi-copy gene analysis for all salivary proteins

Consistent with prior vector genome assemblies, variation within and/or between loci seems to have affected the initial assembly of SP proteins in this species. In general, comparing 2 individual sand fly samples relative to the reference assembly implies that some SP proteins may exist as 2 separate loci given its coverage is significantly less than expected (p <0.0002; see Methods) using the Poisson-based coverage model of Lander and Waterman [46]. The only potential evidence of multiple copies occurred at the 3’ terminal end of the second and third region (exon) of SP42 (2-3X expectation in both samples); however, other regions of the same gene were found less than expected (p < 0.0002) (S2 Table). We conclude that there is no evidence of over assembly of SP proteins in the current Ppap reference assembly.

## Discussion

We examined the genetic variability and potential immunogenicity of nine abundantly expressed *P. papatasi* salivary proteins with the overarching goal of identifying prospective targets to incorporate into an anti-leishmanial vaccine. The salivary proteins assessed included: PpSP12, PpSP14, PpSP28, PpSP29, PpSP30, PpSP32, PpSP36, PpSP42, and PpSP44. All sand flies collected from three natural populations were subjected to similar analyses outlined by Ramalho-Ortigão *et al*. (2015), to ascertain those salivary proteins that demonstrate similar characteristics to PpSP15 as it has been extensively studied as a vaccine target [61]. A multitude of considerations must be addressed to characterize a salivary protein as a potential vaccine candidate, including genetic variability and conservation across populations, consistent expression, and immunogenicity like Th1 vs. Th2 response, and human MHC alleles. We recommend that PpSP12 and PpSP14 be considered in vaccination strategies as these proteins are conserved across populations, demonstrate minimal variability, do not appear to be under selective pressure, and have the potential to activate the human immune system. PpSP28, PpSP29, PpSP32, PpSP36, PpSP42, or PpSP44 may be viable candidates for further vaccination applications but we would prioritize PpSP12 and PpSP14.

PpSP12 and PpSP14, exhibited a high degree of conservation at the nucleotide and amino acid levels (Figs 2 & 5) across all three populations studied. When our sampled sequences were aligned with previously published *P. papatasi* salivary protein gene sequences from Tunisia (*PpSP12* accession number JQ988874 and *PpSP14* accession number JQ988880)[13] and Israel (*PpSP12* accession number AF335485 and *PpSP14* accession number AF335486)[62], *PpSP12* and *PpSP14* demonstrated almost identical sequences with 95% and 91% percent identity shared, respectively (data not shown). This level of conservation across multiple populations beyond those included in this study demonstrates the potential for a vaccine with broad geographic coverage. Furthermore, the population genetics indices do not indicate that PpSP12 is under selective pressure. PpSP14 might be under slight selective pressure as evidenced by Tajima’s D and Ka/Ks ratio though these values are not statistically significant (refer to S39 Table for summary information). In addition, a smaller number of nonsynonymous mutations or replacement changes are observed in PpSP12 (6) and PpSP14 (13) than in previous PpSP15 (19) analysis, further suggesting PpSP12 and PpSP14 are not under positive selection pressures [24]. Nor do the median joining networks utilizing these genes indicate any clear population structuring.

PpSP12, PpSp14, and PpSP15 belong to the family of small odorant-binding proteins (OBP) but their specific functions are unknown. *Phlebotomus* OBPs are related to the D7 protein family that includes PpSP28 and PpSP30 and may have arose from a duplication event of a D7 gene [13]. The high degree of conservancy among the OBPs demonstrated in this study mimics similar conservation of salivary proteins in geographically distant populations of *P. duboscqi* in Kenya and Mali [63]. *P. duboscqi* and *P. papatasi* belong to the same subgenus and are both known vectors of *Le. major*. The use of highly conserved salivary proteins across sand fly species to elicit a cross-protective effect would make the ideal vaccine, and cross protection against *Le. major* using salivary gland homogenate from *P. papatasi* and *P. duboscqi* using a murine model has been demonstrated [17]. Unfortunately, the same cross-protective phenomenon is not observed in phylogenetically distant species like *P. papatasi* and *Lu. longipalpis* [64]. Even though species specificity exists, cross protection may be possible across species that vector the same *Leishmania* parasites, i.e. *Le. major* vectored by *P. papatasi, P. duboscqi,* and *P. bergeroti*. Cross protection might theoretically be possible against sand flies that vector different *Leishmania* species but results in the same clinical disease outcome. For example, cross protection might be possible between *P. papatasi* and *P. sergenti* that vector *Le. major* and *Le. tropica*, respectively. However, this same phenomenon might not be possible with sand flies that vector *Leishmania* parasites that result in different clinical outcomes (i.e. cutaneous and visceral leishmaniasis) [17].

Gene expression is another important consideration in vaccine design as it relates to antigen dosage [65]. Salivary protein genes that are constitutively expressed are viable vaccine targets more so than those genes that change due to seasonality or other factors. Over the course of three sand fly trappings (June, August, September), only *PpSP12* was significantly upregulated in September for the PPJS population but no significant change occurred in the other populations [41]. *PpSP14* did not experience a significant change in expression during the sampling season. Sugar content in plants from dry habitats, like Swaimeh, Jordan, varies in comparison to plants found in irrigated areas like Aswan, Egypt, suggesting that sugar source may be a principle factor in the differential expression demonstrated by *PpSP12*. Gene expression of *PpSP12* and *PpSP14* is influenced by diet and senescence [66]. In colony-reared, 3-day old sand flies, a 3.95 and 2.18-fold change was observed in blood-fed and sugar-fed flies respectively, compared to nonfed sand flies for *PpSP12*. For *PpSP14*, there was a 3.05-fold change in blood fed females compared to nonfed female sand flies. There was similar upregulation of both *PpSP12* and *PpSP14* at day 5 and day 9 post-emergence. Though diet and senescence may influence salivary gland gene expression, environmental factors play a much larger role in gene expression regulation in wild populations [66]. Both *PpSP12* and *PpSP14* were expressed throughout seasonal trappings and when specifically tested for age or diet. Although these proteins are not considered constitutively expressed like *PpSP32*, they do not experience downregulation providing further evidence of their potential to provide a high enough antigen dose to prime the immune system for protection [65,66].

Another key aspect to vaccine development is the potential to elicit an immune response in human hosts. If certain salivary proteins are not predicted to interface with the appropriate human immune cells, then those salivary proteins should be excluded from further study. Both mature PpSP12 and PpSP14 proteins have multiple promiscuous MHC class II epitopes identified for presentation to T-cell receptors with limited variation in the potential epitope regions. Conversely, PpSP28 demonstrates high variability in predicted epitope regions decreasing the bonding likelihood with MHC class II receptors. We also identified the MHC class II alleles expected to recognize the salivary protein epitopes and investigated the predominant alleles of human populations living in Egypt and Jordan. The MHC class II alleles with strong binding affinities for PpSP12 that are also prevalent in Egyptian and Jordanian human populations include: DRB1_0301, DRB1_040X, DRB1_110X, DRB1_1301, and DRB1_150X [67–69]. The MHC class II alleles identified for PpSP14 include: DRB1_1101, DRB1_0301, DRB1_040X, DRB1_11XX, DRB1_13XX, DRB1_15XX, and to a lesser extent DRB1_070X [67–69]. The six remaining salivary proteins are predicted to bind to MHC class II alleles with varying affinity. PpSP36, PpSp42, and PpSP44 demonstrated greater binding affinities to multiple regions for each predicted protein structure but were not predicted to bind with the most prevalent alleles in the human populations from Egypt and Jordan (data not shown). PpSP29, PpSP30, and PpSP32, displayed fewer predicted binding regions with lower affinities for those regions (data not shown). The data from the prediction software tools adds to the mounting evidence in support of using PpSP12 and PpSP14 in vaccination strategies.

Of critical importance is whether these salivary proteins are recognized by human plasma. Although PpSP12 and PpSP14 are smaller in size than the other salivary proteins analyzed, they are less variable overall as there is less opportunity for mutations to occur. Even though larger proteins might be more immunogenic, our data, supported by previous studies, indicate that PpSP12 and PpSP14 will be recognized by alleles circulating in study areas [67–70]. Both PpSP12 and PpSP14 are recognized by the immune system but antibody specificity differs among the human populations tested [70]. Our assessment of human responses included Egyptian and Jordanian residents (MENA donors) and U.S. military personnel deployed overseas. Eleven highly expressed salivary proteins were tested for their antibody specificity when compared to controls (i.e., MENA individuals not living in sand fly endemic regions or U.S. military that have not traveled to *P. papatasi*-endemic regions). MENA donors displayed specificity to PpSP12, PpSP26, PpSP30, PpSP38, and PpSP44 but not PpSP14, whereas U.S. military displayed specificity to PpSP14 and PpSP38 but not PpSP12 [70]. In an independent study, it was shown that plasma antibody specificity of 200 Tunisian children ages 6 to 12 years old reacted to PpSP12, PpSP15, PpSP21, PpSP28, PpSP30, PpSP36, and PpSP44, but not to PpSP14 [19], emphasizing the impact of prolonged exposure to sand fly bites versus naïve individuals traveling to sand fly endemic areas [19,70]. Interestingly, PpSP12 and PpSP14 were also shown immunoreactive in unexposed control donors and that the circulating antibodies against these specific salivary proteins could be the result of exposure to other hematophagous arthropod species [70].

Specific antibody response also factors into the polarization of the immune response to a Th1-mediated or Th2-mediated response. The polarization to Th1 or Th2 responses result in protection against CL or a disease exacerbation effect, respectively [15,62]. In one study, total *P. papatasi* salivary gland homogenate elicited IgG4 specificity as the dominant isotype and subclass circulating in human donors and positively correlated with IgE concentrations [70]. IgG4 and IgE are hallmarks of a Th2 and allergic hypersensitivity response [71]. Another study demonstrated that whole salivary gland homogenate upregulates interleukin 4 (IL-4) while inhibiting interleukin 12 (IL-12) and IFN-γ skewing to a Th2 response in the murine model [72]. Th1/Th2 polarization is also dependent on no exposure or pre-exposure to sand fly bites (as reviewed in [73]). The antibody response to individual salivary protein antigens was characterized [19]. PpSP12 was recognized predominately by IgG1 and IgG2 and not IgG4 nor IgE indicating its potential to polarize to a protective Th1 response. PpSP14 was not characterized as it did not demonstrate antibody specificity, but in another study produced a humoral response [8,19].

Taken together, our results and those of others demonstrate the potential of PpSP12 and PpSp14 as vaccine targets. Further testing needs to be conducted to more specifically determine the Th1/Th2 response of PpSP12 and PpSP14 as well as determine if these proteins would confer protection in individuals living in endemic regions as well as naïve populations who may work or travel to endemic areas. This work, taken together with other studies, indicates that a combinatorial vaccine comprised of specific salivary proteins and a *Leishmania* parasite antigen would confer a more robust immune response resulting in lasting immunity.

## Acknowledgments

We are grateful to the Egyptian Ministry of Health for their aid and in sand fly collections and the Multi National Force and Observers (MFO) military units for transportation in the Sinai Peninsula. Special gratitude goes to Ms. Maria Badra from U.S. Naval Medical Research Unit Number Three (NAMRU-3), Cairo, Egypt, for her organizational skills and support of the work in Egypt. The study protocol was approved by the U.S. Naval Medical Research Unit Number Three (NAMRU-3) Institutional Review Board IRB No. 193, DoD No, NAMRU3.2006.0011, in compliance with all applicable Federal regulations governing the protection of human subjects. The content is solely the responsibility of the authors and does not necessarily represent the official views of the National Institute of Allergy and Infectious Diseases or the National Institutes of Health of the U.S. Department of Defense. DFH is a retired military service member; MRO, IVCA, HAH, SSEH, EEDYF and SK are employees of the U.S. Government. This work was prepared as part of our official duties. Title 17 U.S.C. §105 provides that ’Copyright protection under this title is not available for any work of the United States Government’. Title 17 U.S.C. §101 defines a U.S. Government work as a work prepared by a military service member or employee of the U.S. Government as part of that person’s official duties. The opinions and assertions expressed herein are those of the author(s) and do not necessarily reflect the official policy or position of the Uniformed Services University or the Department of Defense. This work was prepared by a military or civilian employee of the U.S. Government as part of the individual’s official duties and therefore is in the public domain and does not possess copyright protection. The authors thank Mariha Wadsworth, Jonathon Weyerbacher, and Theresa Lai for their help in the sequencing of the sand fly samples.

---

**S1 Table. *Phlebotomus papatasi* salivary protein primers and GenBank accession numbers.**

***=Amplicon is under 200 base pairs; not assigned an accession number; sequences available upon request.

**S2 Table. *Phlebotomus papatasi* salivary protein gene multi-copy assessment.**

**S3 Fig. PpSP29 nucleotide and amino acid variation.**

(A) Weblogo illustrating the relative frequencies of nucleotide polymorphisms in wild caught *P. papatasi* populations from PPAW, PPJM, and PPJS. (B) Weblogo illustrating the relative frequencies of amino acid polymorphisms in wild caught *P. papatasi* populations from PPAW, PPJM, and PPJS.

**S4 Table. PpSP29 population genetics analyses for *P. papatasi* populations**

NS=*p*>0.10; NS^1^=0.10 > *p* > 0.05; *=*p*<0.05

**S5 Table. *PpSP29* pairwise comparisons of genetic differentiation estimates.**

**S6 Fig. Median-joining network for PpSP29 *P. papatasi* haplotypes.**

Circle size and circle color indicates frequency and geographical location of haplotypes, respectively. Haplotype numbers are written next to the corresponding circle H_XX. Red numbers between haplotypes indicate number of mutations between haplotypes.

**S7 Table. PpSP29 sliding window analysis.**

Ka/Ks were plotted for every 70 codons. Values greater than one suggest the potential for positive selection. ----indicates a lack of polymorphic data in the window to calculate a Ka/Ks value.

**S8 Fig. PpSp29 secondary structure, polymorphic sites, and MHC class II epitope predictions.**

The mature PpSP29 amino acid sequence predicted secondary structure. Yellow highlighted amino acids indicate the predicted MHC class II predicted promiscuous peptides. Individual amino acids underlined in black indicate unique polymorphic sites. Predicted secondary structure based on sequence accession #AGE83096[13].

**S9 Fig. PpSP30 nucleotide and amino acid variation.**

(A) Weblogo illustrating the relative frequencies of nucleotide polymorphisms in wild caught *P. papatasi* populations from PPAW, PPJM, and PPJS. (B) Weblogo illustrating the relative frequencies of amino acid polymorphisms in wild caught *P. papatasi* populations from PPAW, PPJM, and PPJS.

**S10 Table. PpSP30 population genetics analyses for *P. papatasi* populations**

NS=*p*>0.10; NS^1^=0.10 > *p* > 0.05; *=*p*<0.05

**S11 Table. *PpSP30* pairwise comparisons of genetic differentiation estimates.**

**S12 Fig. Median-joining network for PpSP30 *P. papatasi* haplotypes.**

Circle size and circle color indicates frequency and geographical location of haplotypes, respectively. Haplotype numbers are written next to the corresponding circle H_XX. Red numbers between haplotypes indicate number of mutations between haplotypes.

**S13 Table. PpSP30 sliding window analysis.**

Ka/Ks were plotted for every 70 codons. Values greater than one suggest the potential for positive selection. ----indicates a lack of polymorphic data in the window to calculate a Ka/Ks value.

**S14 Fig. PpSp30 secondary structure, polymorphic sites, and MHC class II epitope predictions.**

The mature PpSP30 amino acid sequence predicted secondary structure. Yellow highlighted amino acids indicate the predicted MHC class II predicted promiscuous peptides. Individual amino acids underlined in black indicate unique polymorphic sites. Predicted secondary structure based on sequence accession #AGE83093[13].

**S15 Fig. PpSP32 nucleotide and amino acid variation.**

(A) Weblogo illustrating the relative frequencies of nucleotide polymorphisms in wild caught *P. papatasi* populations from PPAW, PPJM, and PPJS. (B) Weblogo illustrating the relative frequencies of amino acid polymorphisms in wild caught *P. papatasi* populations from PPAW, PPJM, and PPJS.

**S16 Table. PpSP32 population genetics analyses for *P. papatasi* populations**

NS=*p*>0.10; NS^1^=0.10 > *p* > 0.05; *=*p*<0.05

**S17 Table. *PpSP32* pairwise comparisons of genetic differentiation estimates.**

**S18 Fig. Median-joining network for PpSP32 *P. papatasi* haplotypes.**

Circle size and circle color indicates frequency and geographical location of haplotypes, respectively. Haplotype numbers are written next to the corresponding circle H_XX. Red numbers between haplotypes indicate number of mutations between haplotypes.

**S19 Table. PpSP32 sliding window analysis.**

Ka/Ks were plotted for every 70 codons. Values greater than one suggest the potential for positive selection. ----indicates a lack of polymorphic data in the window to calculate a Ka/Ks value.

**S20 Fig. PpSp32 secondary structure, polymorphic sites, and MHC class II epitope predictions.**

The mature PpSP32 amino acid sequence predicted secondary structure. Yellow highlighted amino acids indicate the predicted MHC class II predicted promiscuous peptides. Individual amino acids underlined in black indicate unique polymorphic sites. Predicted secondary structure based on sequence accession #AGE83097[13].

**S21 Fig. PpSP36 nucleotide and amino acid variation.**

(A) Weblogo illustrating the relative frequencies of nucleotide polymorphisms in wild caught *P. papatasi* populations from PPAW, PPJM, and PPJS. (B) Weblogo illustrating the relative frequencies of amino acid polymorphisms in wild caught *P. papatasi* populations from PPAW, PPJM, and PPJS.

**S22 Table. PpSP36 population genetics analyses for *P. papatasi* populations**

NS=*p*>0.10; NS^1^=0.10 > *p* > 0.05; *=*p*<0.05

**S23 Table. *PpSP36* pairwise comparisons of genetic differentiation estimates.**

**S24 Fig. Median-joining network for PpSP36 *P. papatasi* haplotypes.**

Circle size and circle color indicates frequency and geographical location of haplotypes, respectively. Haplotype numbers are written next to the corresponding circle H_XX. Red numbers between haplotypes indicate number of mutations between haplotypes.

**S25 Table. PpSP36 sliding window analysis**.

Ka/Ks were plotted for every 70 codons. Values greater than one suggest the potential for positive selection. ----indicates a lack of polymorphic data in the window to calculate a Ka/Ks value.

**S26 Fig. PpSp36 secondary structure, polymorphic sites, and MHC class II epitope predictions.**

The mature PpSP36 amino acid sequence predicted secondary structure. Yellow highlighted amino acids indicate the predicted MHC class II predicted promiscuous peptides. Individual amino acids underlined in black indicate unique polymorphic sites. Predicted secondary structure based on sequence accession #AGE83101[13].

**S27 Fig. PpSP42 nucleotide and amino acid variation.**

(A) Weblogo illustrating the relative frequencies of nucleotide polymorphisms in wild caught *P. papatasi* populations from PPAW, PPJM, and PPJS. (B) Weblogo illustrating the relative frequencies of amino acid polymorphisms in wild caught *P. papatasi* populations from PPAW, PPJM, and PPJS.

**S28 Table. PpSP42 population genetics analyses for *P. papatasi* populations**

NS=*p*>0.10; NS^1^=0.10 > *p* > 0.05; *=*p*<0.05

**S29 Table. *PpSP42* pairwise comparisons of genetic differentiation estimates.**

**S30 Fig. Median-joining network for PpSP42 *P. papatasi* haplotypes.**

Circle size and circle color indicates frequency and geographical location of haplotypes, respectively. Haplotype numbers are written next to the corresponding circle H_XX. Red numbers between haplotypes indicate number of mutations between haplotypes.

**S31 Table. PpSP42 sliding window analysis.**

Ka/Ks were plotted for every 70 codons. Values greater than one suggest the potential for positive selection. ----indicates a lack of polymorphic data in the window to calculate a Ka/Ks value.

**S32 Fig. PpSp42 secondary structure, polymorphic sites, and MHC class II epitope predictions.**

The mature PpSP42 amino acid sequence predicted secondary structure. Yellow highlighted amino acids indicate the predicted MHC class II predicted promiscuous peptides. Individual amino acids underlined in black indicate unique polymorphic sites. Predicted secondary structure based on sequence accession #AGE83094[13].

**S33 Fig. PpSP44 nucleotide and amino acid variation.**

(A) Weblogo illustrating the relative frequencies of nucleotide polymorphisms in wild caught *P. papatasi* populations from PPAW, PPJM, and PPJS. (B) Weblogo illustrating the relative frequencies of amino acid polymorphisms in wild caught *P. papatasi* populations from PPAW, PPJM, and PPJS.

**S34 Table. PpSP44 population genetics analyses for *P. papatasi* populations**

NS=*p*>0.10; NS^1^=0.10 > *p* > 0.05; *=*p*<0.05

**S35 Table. *PpSP44* pairwise comparisons of genetic differentiation estimates.**

**S36 Fig. Median-joining network for PpSP44 *P. papatasi* haplotypes.**

Circle size and circle color indicates frequency and geographical location of haplotypes, respectively. Haplotype numbers are written next to the corresponding circle H_XX. Red numbers between haplotypes indicate number of mutations between haplotypes.

**S37 Table. PpSP44 sliding window analysis.**

Ka/Ks were plotted for every 70 codons. Values greater than one suggest the potential for positive selection. ----indicates a lack of polymorphic data in the window to calculate a Ka/Ks value.

**S38 Fig. PpSp44 secondary structure, polymorphic sites, and MHC class II epitope predictions.**

The mature PpSP44 amino acid sequence predicted secondary structure. Yellow highlighted amino acids indicate the predicted MHC class II predicted promiscuous peptides. Individual amino acids underlined in black indicate unique polymorphic sites. Predicted secondary structure based on sequence accession #AGE83095[13].

**S39 Table. Summary Tajima’s D and Ka/Ks analysis for all *P. papatasi* salivary proteins studied.**

NS=*p*>0.10; NS^1^=0.10 > *p* > 0.05; *=*p*<0.05

## References

1. Alvar J, Vélez ID, Bern C, Herrero M, Desjeux P, Cano J, et al. Leishmaniasis worldwide and global estimates of its incidence. PLoS One. 2012;7. doi:10.1371/journal.pone.0035671

2. World Health Organization. Control of the leishmaniases. World Heal Organ Tech Rep Ser. Geneva; 2010;949: 1–186. doi:10.1038/nrmicro1766

3. Ribeiro JM. Role of saliva in blood feeding by arthropods. Ann Rev Entomol. 1987;32: 463–478. doi:10.1146/annurev.en.32.010187.002335

4. Marzouki S, Abdeladhim M, Abdessalem C Ben, Oliveira F, Ferjani B, Gilmore D, et al. Salivary antigen SP32 is the immunodominant target of the antibody response to *Phlebotomus papatasi* bites in humans. PLoS Negl Trop Dis. 2012;6. doi:10.1371/journal.pntd.0001911

5. Sima M, Novotny M, Pravda L, Sumova P, Rohousova I, Volf P. The diversity of yellow-related proteins in sand flies (Diptera: Psychodidae). Traub-Csekö YM, editor. PLoS One. 2016;11: e0166191. doi:10.1371/journal.pone.0166191

6. Marzouki S, Kammoun-Rebai W, Bettaieb J, Abdeladhim M, Hadj Kacem S, Abdelkader R, et al. Validation of recombinant salivary protein PpSP32 as a suitable marker of human exposure to *Phlebotomus papatasi*, the vector of *Leishmania major* in Tunisia. PLoS Negl Trop Dis. 2015;9: 1–14. doi:10.1371/journal.pntd.0003991

7. Valenzuela JG, Garfield M, Rowton ED, Pham VM. Identification of the most abundant secreted proteins from the salivary glands of the sand fly *Lutzomyia longipalpis*, vector of *Leishmania chagasi*. J Exp Biol. 2004;207: 3717–3729. doi:10.1242/jeb.01185

8. Oliveira F, Lawyer PG, Kamhawi S, Valenzuela JG. Immunity to distinct sand fly salivary proteins primes the anti-leishmania immune response towards protection or exacerbation of disease. PLoS Negl Trop Dis. 2008;2. doi:10.1371/journal.pntd.0000226

9. Oliveira F, Jochim RC, Valenzuela JG, Kamhawi S. S and flies, *Leishmania*, and transcriptome-borne solutions. Parasitol Int. Elsevier B.V.; 2009;58: 1–5. doi:10.1016/j.parint.2008.07.004

10. Oliveira F, Rowton E, Aslan H, Gomes R, Castrovinci PA, Alvarenga PH, et al. A sand fly salivary protein vaccine shows efficacy against vector-transmitted cutaneous leishmaniasis in nonhuman primates. Sci Transl Med. 2015;7: 290ra90. doi:10.1126/scitranslmed.aaa3043

11. Tlili A, Marzouki S, Chabaane E, Abdeladhim M, Kammoun-Rebai W, Sakkouhi R, et al. *Phlebotomus papatasi* yellow-related and apyrase salivary proteins are candidates for vaccination against human cutaneous leishmaniasis. J Invest Dermatol. Elsevier; 2017;138: 598–606. doi:10.1016/J.JID.2017.09.043

12. Kamhawi S, Belkaid Y, Modi G, Rowton E, Sacks D. Protection against cutaneous leishmaniasis resulting from bites of uninfected sand flies. Science. 2000;290: 1351–1354. doi:10.1126/science.290.5495.1351

13. Abdeladhim M, Jochim RC, Ben Ahmed M, Zhioua E, Chelbi I, Cherni S, et al. Updating the salivary gland transcriptome of *Phlebotomus papatasi* (Tunisian strain): The search for sand fly-secreted immunogenic proteins for humans. PLoS One. 2012;7: e47347. doi:10.1371/journal.pone.0047347

14. Titus R, Ribeiro J. Salivary gland lysates from the sand fly *Lutzomyia longipalpis* enhance *Leishmania* infectivity. Science (80-). 1988;239: 1306–1308. doi:10.1126/science.3344436

15. Belkaid Y, Kamhawi S, Modi G, Valenzuela J, Noben-Trauth N, Rowton E, et al. Development of a natural model of cutaneous leishmaniasis: Powerful effects of vector saliva and saliva preexposure on the long-term outcome of *Leishmania major* infection in the mouse ear dermis. J Exp Med. 1998;188: 1941–1953. doi:10.1084/jem.188.10.1941

16. Rohousova I, Ozensoy S, Ozbel Y, Volf P. Detection of species-specific antibody response of humans and mice bitten by sand flies. Parasitology. 2005;130: 493–499. doi:10.1017/S003118200400681X

17. Lestinova T, Vlkova M, Votypka J, Volf P, Rohousova I. *Phlebotomus papatasi* exposure cross-protects mice against *Leishmania major* co-inoculated with *Phlebotomus duboscqi* salivary gland homogenate. Acta Trop. 2015;144: 9–18. doi:10.1016/j.actatropica.2015.01.005

18. Tavares NM, Silva RA, Costa DJ, Pitombo MA, Fukutani KF, Miranda JC, et al. *Lutzomyia longipalpis* saliva or salivary protein LJM19 protects against *Leishmania braziliensis* and the saliva of its vector, *Lutzomyia intermedia*. PLoS Negl Trop Dis. 2011;5. doi:10.1371/journal.pntd.0001169

19. Marzouki S, Ben Ahmed M, Boussoffara T, Abdeladhim M, Ben Aleya-Bouafif N, Namane A, et al. Characterization of the antibody response to the saliva of *Phlebotomus papatasi* in people living in endemic areas of cutaneous leishmaniasis. Am J Trop Med Hyg. 2011;84: 653–661. doi:10.4269/ajtmh.2011.10-0598

20. de Moura TR, Oliveira F, Novais FO, Miranda JC, Clarêncio J, Follador I, et al. Enhanced *Leishmania braziliensis* infection following pre-exposure to sandfly saliva. PLoS Negl Trop Dis. 2007;1. doi:10.1371/journal.pntd.0000084

21. Andrade BB, Teixeira CR. Biomarkers for exposure to sand flies bites as tools to aid control of leishmaniasis. Front Immunol. 2012;3: 1–7. doi:10.3389/fimmu.2012.00121

22. Mondragon-Shem K, Al-Salem WS, Kelly-Hope L, Abdeladhim M, Al-Zahrani MH, Valenzuela JG, et al. Severity of Old World cutaneous leishmaniasis is influenced by previous exposure to sandfly bites in Saudi Arabia. PLoS Negl Trop Dis. 2015;9: e0003449. doi:10.1371/journal.pntd.0003449

23. Elnaiem D-EA, Meneses C, Slotman M, Lanzaro GC. Genetic variation in the sand fly salivary protein, SP-15, a potential vaccine candidate against Leishmania major. Insect Mol Biol. 2005;14: 145–150. doi:10.1111/j.1365-2583.2004.00539.x

24. Ramalho-Ortigão M, Coutinho-Abreu I V., Balbino VQ, Figueiredo CAS, Mukbel R, Dayem H, et al. *Phlebotomus papatasi* SP15: mRNA expression variability and amino acid sequence polymorphisms of field populations. Parasit Vectors. 2015;8: 298. doi:10.1186/s13071-015-0914-2

25. Coutinho-Abreu I V., Ramalho-Ortigao M. Ecological genomics of sand fly salivary gland genes: An overview. J Vector Ecol. 2011;36: 58–63. doi:10.1111/j.1948-7134.2011.00112.x

26. Hamarsheh O, Presber W, Abdeen Z, Sawalha S, Al-Lahem A, Schönian G. Genetic structure of Mediterranean populations of the sandfly *Phlebotomus papatasi* by mitochondrial cytochrome b haplotype analysis. Med Vet Entomol. 2007;21: 270–277. doi:10.1111/j.1365-2915.2007.00695.x

27. Hamarsheh O, Presber W, Yaghoobi-Ershadi MR, Amro A, Al-Jawabreh A, Sawalha S, et al. Population structure and geographical subdivision of the *Leishmania major* vector *Phlebotomus papatasi* as revealed by microsatellite variation. Med Vet Entomol. 2009;23: 69–77. doi:10.1111/j.1365-2915.2008.00784.x

28. Depaquit J, Lienard E, Verzeaux-Griffon A, Ferté H, Bounamous A, Gantier JC, et al. Molecular homogeneity in diverse geographical populations of *Phlebotomus papatasi* (Diptera, Psychodidae) inferred from ND4 mtDNA and ITS2 rDNA. Epidemiological consequences. Infect Genet Evol. 2008;8: 159–170. doi:10.1016/j.meegid.2007.12.001

29. Esseghir S, Ready PD, Killick-Kendrick R, Ben-Ismail R. Mitochondrial haplotypes and phylogeography of *Phlebotomus* vectors of *Leishmania major*. Insect Mol Biol. 1997;6: 211–225. doi:DOI 10.1046/j.1365-2583.1997.00175.x

30. Raja B, Jaouadi K, Haouas N, Mezhoud H, Bdira S, Amor S. Mitochondrial cytochrome b variation in populations of the cutaneous leishmaniasis vector *Phlebotomus papatasi* across eastern Tunisia. Int J Biodivers Conserv. 2012;4: 189–196. doi:10.5897/IJBC11.191

31. Flanley CM, Ramalho-Ortigao M, Coutinho-Abreu I V, Mukbel R, Hanafi HA, El-Hossary SS, et al. Population genetics analysis of *Phlebotomus papatasi* sand flies from Egypt and Jordan based on mitochondrial cytochrome b haplotypes. Parasit Vectors. 2018;11: 1–11. doi:10.1186/s13071-018-2785-9

32. Bessat M, Shanat S El. Leishmaniasis: Epidemiology, control and future perspectives with special emphasis on Egypt. J Trop Dis. 2015;2: 1–10. doi:10.4172/2329891X.1000153

33. Salam N, Al-Shaqha WM, Azzi A. Leishmaniasis in the Middle East: Incidence and epidemiology. PLoS Negl Trop Dis. 2014;8: 1–8. doi:10.1371/journal.pntd.0003208

34. Lane RP. The sandflies of Egypt (Diptera: Phlebotominae). Bull Br Museum (Natural Hist. London: The Museum; 1986;52: 1–35.

35. Añez N, Tang Y. Comparison of three methods for age-grading of female Neotropical phlebotomine sandflies. Med Vet Entomol. 1997;11: 3–7. doi:10.1111/j.1365- 2915.1997.tb00283.x

36. STyx. World Location Map. In: Wikimedia Commons [Internet]. 2010. Available: https://commons.wikimedia.org/wiki/File:World_location_map.svg

37. Hoel DF, Butler JF, Fawaz EY, Watany N, El-Hossary SS, Villinski J. Response of phlebotomine sand flies to light-emitting diode-modified light traps in southern Egypt. J Vector Ecol. 2007;32: 302–8. doi:10.3376/1081-1710(2007)32[302:ROPSFT]2.0.CO;2

38. Saliba EK, Oumeish OY. Reservoir hosts of cutaneous leishmaniasis. Clin Dermatol. 1999;17: 275–277. doi:10.1016/S0738-081X(99)00045-0

39. Schlein Y, Jacobson RL. Linkage between susceptibility of *Phlebotomus papatasi* to *Leishmania major* and hunger tolerance. Parasitology. 2002;125: 343–8. doi:10.1017/S0031182002002147

40. Kamhawi S, Modi GB, Pimenta PF, Rowton E, Sacks DL. The vectorial competence of *Phlebotomus sergenti* is specific for *Leishmania tropica* and is controlled by species-specific, lipophosphoglycan-mediated midgut attachment. Parasitology. 2000;121: 25–33. doi:10.1017/S0031182099006125

41. Coutinho-Abreu I V, Mukbel R, Hanafi HA, Fawaz EY, El-Hossary SS, Wadsworth M, et al. Expression plasticity of *Phlebotomus papatasi* salivary gland genes in distinct ecotopes through the sand fly season. BMC Ecol. BioMed Central Ltd; 2011;11: 24. doi:10.1186/1472-6785-11-24

42. Kumar S, Stecher G, Tamura K. MEGA7: Molecular evolutionary genetics analysis version 7.0 for bigger datasets. Mol Biol Evol. 2016;33: 1870–1874. doi:10.1093/molbev/msw054

43. Miles A, Iqbal Z, Vauterin P, Pearson R, Campino S, Theron M, et al. Indels, structural variation, and recombination drive genomic diversity in *Plasmodium falciparum*. Genome Res. 2016;26: 1288–1299. doi:10.1101/gr.203711.115

44. Andrews S. FastQC: a quality control tool for high throughput sequence data. [Internet]. 2010. Available: https://www.bioinformatics.babraham.ac.uk/projects/fastqc/

45. Li H, Durbin R. Fast and accurate long-read alignment with Burrows-Wheeler transform. Bioinformatics. 2010;26: 589–595. doi:10.1093/bioinformatics/btp698

46. Lander ES, Waterman MS. Genomic mapping by fingerprinting random clones: A mathematical analysis. Genomics. 1988;2: 231–239. doi:10.1016/0888-7543(88)90007-9

47. Rozas J, Ferrer-Mata A, Sanchez-DelBarrio JC, Guirao-Rico S, Librado P, Ramos-Onsins SE, et al. DnaSP 6: DNA sequence polymorphism analysis of large data sets. Mol Biol Evol. Oxford University Press; 2017;34: 3299–3302. doi:10.1093/molbev/msx248

48. Hudson RR, Slatkin M, Maddison WP. Estimation of levels of gene flow from DNA sequence data. Genetics. 1992;132: 583–589. doi:PMC1205159

49. Wright S. Evolution and the genetics of populations, variability within and among natural populations. The University of Chicago Press, Chicago. 1978.

50. Nei M. Analysis of gene diversity in subdivided populations. Proc Natl Acad Sci U S A. 1973;70: 3321–3323. doi:10.1073/pnas.70.12.3321

51. Hudson RR, Boos DD, Kaplan NL. A statistical test for detecting geographic subdivision. Mol Biol Evol. 1992;9: 138–151.

52. Kimura M. Evolutionary rate at the molecular level. Nature. 1968;217: 624–626. doi:10.1038/217624a0

53. Tajima F. Statistical method for testing the neutral mutation hypothesis by DNA polymorphism. Genetics. 1989;123: 585–595. doi:PMC1203831

54. Fu YX, Li WH. Statistical tests of neutrality of mutations. Genetics. 1993;133: 693–709.

55. Crooks G, Hon G, Chandonia J, Brenner S. WebLogo: a sequence logo generator. Genome Res. 2004;14: 1188–1190. doi:10.1101/gr.849004.1

56. Bandelt HJ, Forster P, Röhl A. Median-joining networks for inferring intraspecific phylogenies. Mol Biol Evol. 1999;16: 37–48. doi:10.1093/oxfordjournals.molbev.a026036

57. Wang P, Sidney J, Dow C, Mothé B, Sette A, Peters B. A systematic assessment of MHC class II peptide binding predictions and evaluation of a consensus approach. PLoS Comput Biol. 2008;4. doi:10.1371/journal.pcbi.1000048

58. Wang P, Sidney J, Kim Y, Sette A, Lund O, Nielsen M, et al. Peptide binding predictions for HLA DR, DP and DQ molecules. BMC Bioinformatics. 2010;11: 568–580. doi:10.1186/1471-2105-11-568

59. Singh H, Raghava GPS. ProPred: Prediction of HLA-DR binding sites. Bioinformatics. 2001;17: 1236–1237. doi:10.1093/bioinformatics/17.12.1236

60. Betts MJ, Russell RB. Amino acid properties and consequences of substitutions. In: Barnes MR, Gray IC, editors. Bioinformatics for Geneticists. West Sussex: John Wiley & Sons Ltd.; 2003. pp. 289–304. doi:10.1002/0470867302.ch14

61. Srivastava S, Shankar P, Mishra J, Singh S. Possibilities and challenges for developing a successful vaccine for leishmaniasis. Parasites and Vectors. 2016;9: 1–15. doi:10.1186/s13071-016-1553-y

62. Valenzuela JG, Belkaid Y, Garfield MK, Mendez S, Kamhawi S, Rowton ED, et al. Toward a defined anti-*Leishmania* vaccine targeting vector antigens: Characterization of a protective salivary protein. J Exp Med. 2001;194: 331–342. doi:10.1084/jem.194.3.331

63. Kato H, Anderson JM, Kamhawi S, Oliveira F, Lawyer PG, Pham VM, et al. High degree of conservancy among secreted salivary gland proteins from two geographically distant *Phlebotomus duboscqi* sandflies populations (Mali and Kenya). BMC Genomics. 2006;7: 1–21. doi:10.1186/1471-2164-7-226

64. Thiakaki M, Rohousova I, Volfova V, Volf P, Chang KP, Soteriadou K. Sand fly specificity of saliva-mediated protective immunity in *Leishmania amazonensis*-BALB/c mouse model. Microbes Infect. 2005;7: 760–766. doi:10.1016/j.micinf.2005.01.013

65. Henrickson SE, Mempel TR, Mazo IB, Liu B, Artyomov MN, Zheng H, et al. T cell sensing of antigen dose governs interactive behavior with dendritic cells and sets a threshold for T cell activation. Nat Immunol. 2008;9: 282–291. doi:10.1038/ni1559

66. Coutinho-Abreu I V, Wadsworth M, Stayback G, Ramalho-Ortigao M, McDowell MA. Differential expression of salivary gland genes in the female sand fly P*hlebotomus papatasi* (Diptera: Psychodidae). J Med Entomol. 2010;47: 1146–1155. doi:10.1603/ME10072

67. Hajjej A, Almawi WY, Arnaiz-Villena A, Hattab L, Hmida S. The genetic heterogeneity of Arab populations as inferred from HLA genes. PLoS One. 2018;13: 1–24. doi:10.1371/journal.pone.0192269

68. González-Galarza FF, Takeshita LYC, Santos EJM, Kempson F, Maia MHT, da Silva ALS, et al. Allele frequency net 2015 update: New features for HLA epitopes, KIR and disease and HLA adverse drug reaction associations. Nucleic Acids Res. 2015;43: D784– D788. doi:10.1093/nar/gku1166

69. Elbjeirami WM, Abdel-Rahman F, Hussein AA. Probability of finding an HLA-matched donor in immediate and extended families: The Jordanian experience. Biol Blood Marrow Transplant. 2013;19: 221–226. doi:10.1016/j.bbmt.2012.09.009

70. Geraci NS, Mukbel RM, Kemp MT, Wadsworth MN, Lesho E, Stayback GM, et al. Profiling of human acquired immunity against the salivary proteins of *Phlebotomus papatasi* reveals clusters of differential immunoreactivity. Am J Trop Med Hyg. 2014;90: 923–938. doi:10.4269/ajtmh.13-0130

71. Punnonen J, Aversa G, Cocks BG, McKenzie AN, Menon S, Zurawski G, et al. Interleukin 13 induces interleukin 4-independent IgG4 and IgE synthesis and CD23 expression by human B cells. Proc Natl Acad Sci. 1993;90: 3730–3734. doi:10.1073/pnas.90.8.3730

72. Mbow ML, Bleyenberg JA, Hall LR, Titus RG. *Phlebotomus papatasi* sand fly salivary gland lysate down-regulates a Th1, but up-regulates a Th2, response in mice infected with *Leishmania major*. J Immunol. 1998;161: 5571–7.

73. Lestinova T, Rohousova I, Sima M, de Oliveira CI, Volf P. Insights into the sand fly saliva: Blood-feeding and immune interactions between sand flies, hosts, and *Leishmania*. PLoS Negl Trop Dis. 2017;11: 1–26. doi:10.1371/journal.pntd.0005600

